# Tolerance thresholds underlie responses to DNA damage during germline development

**DOI:** 10.1101/2024.01.07.574510

**Authors:** Gloria Jansen, Daniel Gebert, Tharini Ravindra Kumar, Emily Simmons, Sarah Murphy, Felipe Karam Teixeira

## Abstract

Selfish DNA modules like transposable elements (TEs) are particularly active in the germline, the lineage that passes genetic information across generations. New TE insertions can disrupt genes and impair the functionality and viability of germ cells. However, we find that in *P*-*M* hybrid dysgenesis in *Drosophila*, a sterility syndrome triggered by the *P*-element DNA transposon, germ cells harbour unexpectedly few new TE insertions, despite accumulating DNA double-strand breaks (DSBs) and inducing cell cycle arrest. Using an engineered CRISPR-Cas9 system, we show that generating DSBs at silenced *P*-elements or other non-coding sequences is sufficient to induce germ cell loss independently of gene disruption. Indeed, we demonstrate that both developing and adult mitotic germ cells are sensitive to DSBs in a dosage-dependent manner. Following the mitotic-to-meiotic transition, however, germ cells become more tolerant to DSBs, completing oogenesis regardless of the accumulated genome damage. Our findings establish DNA damage tolerance thresholds as crucial safeguards of genome integrity during germline development.

## Introduction

The germline is the cell lineage that is responsible for the inheritance of genetic information in multicellular eukaryotes. In most animal species, germ cells are set aside from somatic lineages during early embryonic development and follow a unique developmental programme that culminates in the production of haploid gametes carrying the genetic material that is passed to the next generation (Dansereau and Lasko, 2008). Due to its central role in genetic inheritance, the germline is known to be the battleground where genetic conflicts between selfish genetic elements, such as TEs, and the host genome take place (Cosby et al., 2019; Doolittle and Sapienza, 1980; Orgel and Crick, 1980). Excessive TE activity fulfils the selfish drive of TEs to increase in copy number per genome but can impair genome integrity and functioning, thereby threatening the faithful transmission of genetic information (Bourque et al., 2018; Haig, 2016). Unsurprisingly, several host mechanisms exist to limit the activity of TEs in the germline, minimising changes in the inherited genetic material (Ecco et al., 2017; Malone et al., 2009; Molaro and Malik, 2016).

A textbook example of the detrimental effects of uncontrolled TE activity on the germline is provided by the *P*-*M* hybrid dysgenesis system in *Drosophila melanogaster.* In this system, excessive activity of the *P*-element transposon triggers chromosomal rearrangements and increased mutation rates, ultimately leading to sterility (Bingham et al., 1982; Kidwell et al., 1977; Kidwell and Novy, 1979). This syndrome specifically affects the germline of the progeny arising from crosses between *P*-element-containing (*P*) and *P*-element-devoid (*M*) strains in a non-reciprocal manner (Bingham et al., 1982; Kidwell et al., 1977). In the progeny of females from *M-*strains crossed to males from *P*-strains, germ cells are lost during development, leading to fully sterile adults (known as dysgenic hybrids) (Bingham et al., 1982; Kidwell and Novy, 1979; Teixeira et al., 2017). Progeny produced from the reciprocal cross, between *P*-strain females and *M-*strain males (known as non-dysgenic hybrids), are protected from *P*-element activity by a maternally inherited, small RNA-based TE silencing mechanism, and therefore are fully fertile (Aravin et al., 2007; Brennecke et al., 2008; Teixeira et al., 2017).

The *P*-element is a DNA transposon, a class of TEs that transpose by cut-and-paste mechanisms (Bingham et al., 1982; Engels et al., 1990; Kaufman and Rio, 1992). Full-length *P*-elements (2.9-kilobases) encode a single polypeptide, *P*-transposase, which recognises specific DNA sequence motifs just inside of the *P*-element 5’ and 3’ terminal inverted repeats (TIRs) (Kaufman et al., 1989). Upon binding, the *P*-transposase cleaves both DNA strands in a defined manner, excising the target DNA to form the strand transfer complex (Beall and Rio, 1997; Kaufman and Rio, 1992; Rio and Rubin, 1988). This complex then mediates the integration of the excised DNA element at a different genomic locus, completing the transposition process (Beall and Rio, 1998). Consequently, *P*-element transposition leads to sequence changes at both the excision and insertion loci (Bellen et al., 2011; Bellen et al., 2004).

The prevailing model for how *P*-element activity leads to germ cell loss during dysgenesis postulates that mutagenesis and gene disruption, caused by new *P*-element insertions into coding regions, ultimately impairs cell functioning and viability (Griffiths et al., 2000; Khurana et al., 2011). However, given that the diploid *Drosophila* genome is largely haplosufficient (Lindsley et al., 1972), it has been challenging to reconcile how new *P*-element insertions, which affect only one of two alleles (Engels et al., 1990), could reproducibly affect germ cell function. Moreover, since *P*-elements are known to preferentially insert into non-coding regions, including promoters and regions overlapping origins of replication (Spradling et al., 2011), it remains unclear how random new insertions can lead to highly penetrant, lethal mutations. The fully penetrant germ cell death phenotype imposes challenges to precisely determine the rates and the genomic sites of new *P*-element insertions in dysgenic germ cells. As such, this model remains mostly untested.

Despite considerable work dissecting the host silencing pathways controlling TE expression in the germline (Ecco et al., 2017; Malone et al., 2009; Molaro and Malik, 2016), the impact of active TEs that have evaded silencing can have on germ cells remains poorly understood. Here, using *P*-*M* hybrid dysgenesis as a model, we show that excessive TE activity in embryonic germ cells leads to the accumulation of DSBs and persistent cell cycle arrest prior to the fully penetrant germ cell loss phenotype observed in early larval stages. Using FACS sorting coupled with single-cell, whole-genome DNA-sequencing, we find that dysgenic embryonic germ cells acquire surprisingly few new *P*-element insertions, and, in this aspect, are indistinguishable from non-dysgenic PGCs. Given this, we tested whether inducing DNA damage at endogenously silenced *P*-elements, which mimics the excision step of *P*-element transposition, can by itself elicit germ cell loss. Using an engineered, Cas9-based transgenic system to inflict dosage- and sequence-specific DSBs at *P*-elements or at other non-coding sequences of the genome, we demonstrate that embryonic germ cells, as well as mitotically dividing adult germ cells, are sensitive to DSBs in a dosage-dependent manner. By contrast, once germ cells have completed mitotic cycles and acquired programmed DSBs during meiotic recombination, they become tolerant to DSBs, with oogenesis proceeding despite the accumulation of high levels of DNA damage. Taken together, our findings suggest that pre-defined DNA damage tolerance thresholds in the developing germline form a selective barrier that can shape transposon proliferation strategies.

## Results

### PGCs accumulate DSBs and undergo sustained cell cycle arrest during *P*-*M* hybrid dysgenesis

In dysgenic hybrids, germline development proceeds normally during embryogenesis, but PGC numbers decrease from the first instar larval stage (Teixeira et al., 2017). To investigate the levels of DNA damage in dysgenic and non-dysgenic PGCs prior to germ cell loss, we performed antibody staining against the phosphorylated histone H2A variant (pH2Av), a readout of DSBs (Madigan et al., 2002), at successive stages of embryonic development (Figures 1A-C). At the pole cell stage, ∼20-30% of both dysgenic and non-dysgenic PGCs were positive for pH2Av. However, following germline zygotic genome activation (ZGA) at approximately 4 hours after egg laying (Van Doren et al., 1998), an increasing number of dysgenic PGCs accumulated strong pH2Av signal, with around 70% of dysgenic PGCs positive for pH2Av at the early migration stage and over 95% at gonadal coalescence. In contrast, the proportion of non-dysgenic PGCs showing strong pH2Av signal stayed constant at ∼20-30% during these stages.

**Figure 1.**
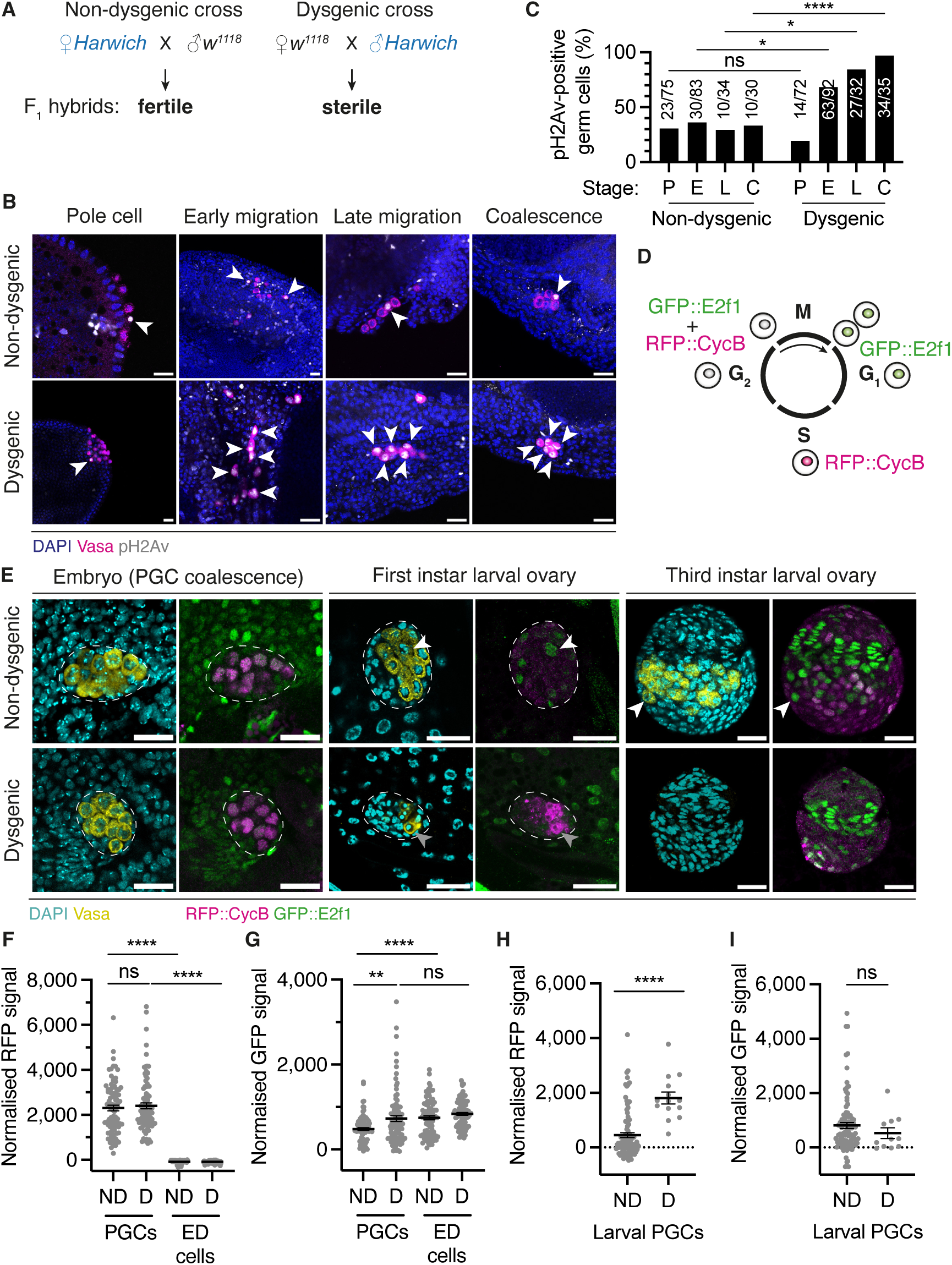
PGCs accumulate DSBs and undergo sustained cell cycle arrest during *P-M* hybrid dysgenesis. **A.** *P*-*M* hybrid dysgenesis crossing scheme. **B.** Dysgenic and non-dysgenic hybrid progeny at four successive stages of embryonic germline development, labelled with DAPI (nuclei, blue), Vasa (germline, magenta) and pH2Av (DSBs, grayscale). White arrowheads indicate pH2Av-positive PGCs. **C.** Proportion of pH2Av-positive PGCs in dysgenic and non-dysgenic hybrid embryos at each developmental stage (P: pole cell stage, E: early migration stage, L: late migration stage and C: coalescence stage). Absolute PGC numbers are shown above bars. **D.** Schematic showing Fly-FUCCI fluorescent readout at each cell cycle stage. **E.** Dysgenic and non-dysgenic hybrid progeny expressing Fly-FUCCI at three developmental stages (embryonic PGC coalescence, first and third instar larva), labelled with DAPI (cyan) and Vasa (yellow) (left hand panels), and RFP (RFP::CycB marker, magenta) and GFP (GFP::E2f1 marker, green) (right hand panels). Dashed lines outline gonads within embryonic or larval tissue. PGCs that re-entered the cell cycle (white arrowheads) or remained arrested (grey arrowheads) are indicated. **F-I.** Quantitation of RFP and GFP fluorescent signal in embryonic PGCs and epidermal (ED) cells (F and G) and first instar larval PGCs (H and I) shown in (E), normalised by signal in somatic epithelial (non-ED) cells. Each data point represents one PGC or ED cell in dysgenic (D) or non-dysgenic (ND) progeny. Black line is mean, error bars are ± SEM. **** p ≤ 0.0001, ** p ≤ 0.01, * p < 0.05 and ns p > 0.05, unpaired *t*-tests. Scale bars, 20 µm.

Cell cycle arrest is a conserved response to genome damage (Melo and Toczyski, 2002; Shim et al., 2014; Xu et al., 2001). To investigate cell cycle dynamics of PGCs during hybrid dysgenesis, we introduced the Fly-FUCCI cell cycle indicator system into the *P*-element-containing (*Harwich*) and *P*-element-devoid (*w^1118^* or *‘white’*) fly strains (Figure 1D) (Zielke et al., 2014). Fly-FUCCI relies on fluorescently tagged, ubiquitously expressed protein reporters whose activity provides a fluorescent readout of the distinct phases of the cell cycle *in vivo.* By crossing these strains reciprocally, we produced dysgenic and non-dysgenic progeny, and used confocal microscopy to determine the cell cycle phase of individual PGCs during embryonic and early larval development. In wild-type, PGCs are known to exit the cell cycle upon their formation at the posterior pole of the embryo and remain stalled in G2-phase for a period of 16-18 hours (Su et al., 1998). A few hours before larval hatching, PGCs progressively re-enter the cell cycle, with robust cycling activity only being observed during larval stages (Williamson and Lehmann, 1996). Quantification of RFP (RFP::CycB; S/G2-phase) and GFP (GFP::E2f1; M/G1/G2-phase) signal in PGCs relative to somatic epidermal cells, which are known to be in G1-phase (Knoblich et al., 1994), indicated that both non-dysgenic and dysgenic PGCs were arrested in G2-phase prior to embryonic gonadal coalescence (Figures 1E-1G). However, while non-dysgenic PGCs re-entered the cell cycle in early larval stages as shown by the loss of RFP::CycB signal, marking their asynchronous transition into M/G1-phase, all dysgenic PGCs remained arrested in G2-phase and failed to re-enter the cell cycle (Figures 1E, 1H and 1I). Together, these analyses indicate that DSBs induced following ZGA are associated with sustained cell cycle arrest and failure to re-enter the cell cycle, and thus represent the earliest signatures of PGC loss during dysgenesis.

### PGCs acquire few new *P*-element insertions during hybrid dysgenesis

In dysgenic PGCs, activation of paternally inherited *P*-elements is hypothesised to induce high numbers of new *P*-element insertions (Griffiths et al., 2000; Khurana et al., 2011; Moon et al., 2018). To precisely determine the number and insertion sites of new *P*-element insertions in individual germ cells during embryogenesis, we isolated GFP-labelled PGCs from dysgenic and non-dysgenic embryos by FACS and performed single-cell whole-genome DNA sequencing analysis (Figure 2A). We isolated PGCs at late embryonic stages, which precedes dysgenic germ cell death and where over 95% of dysgenic PGCs showed strong pH2Av signal and cell cycle arrest (Figures 1B and 1C). Genomic DNA was extracted from individually sorted PGCs and amplified using the PCR-free, isothermal multiple displacement amplification technique (Dean et al., 2001). This procedure generated ∼5-23 μg of Whole-Genome Amplified (WGA) DNA per sorted cell, with an average amplicon size of 10-18 kilobases (Figure S1A). Female and male sorted PGCs were distinguished using quantitative PCR to screen for the presence of the Y chromosome (Figure S1B). The WGA DNA of 40 individual PGCs was then used to generate DNA-sequencing libraries (average genome coverage = 67X) (Figure S1C). These comprised 20 dysgenic and 20 non-dysgenic PGCs, half of which were female and the other half male.

**Figure 2.**
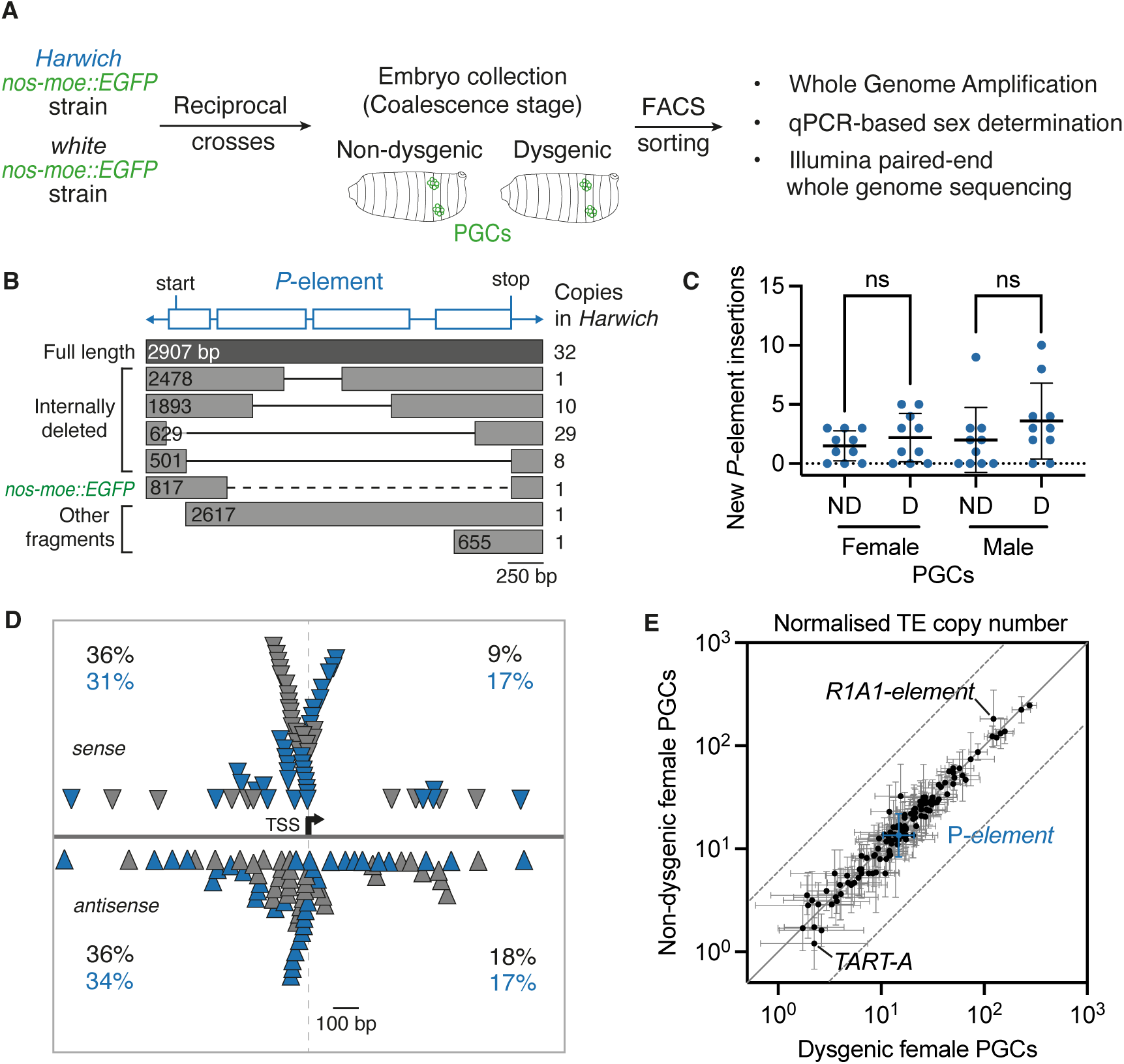
PGCs acquire few new *P*-element insertions during hybrid dysgenesis. **A.** Experimental design for FACS sorting and whole-genome sequencing of single PGCs from dysgenic and non-dysgenic hybrid embryos expressing the transgenic germline marker *nos-moe::EGFP*. **B.** Genomic copy numbers and structures of *P*-elements present in the parental *Harwich* strain, relative to the consensus sequence (blue, with position of start and stop codons indicated). Black solid lines indicate internally deleted sequences. The *nos-moe::EGFP* transgene (dashed line) is flanked by *P*-element sequence. **C.** Quantitation of new *P*-element insertions present in sorted PGCs but not present in *Harwich*. ns p > 0.05, unpaired *t*-test. **D.** Map of *P*-element insertions existing in *Harwich* (grey triangles) and new insertions in PGCs (blue triangles) within 1 kb of the nearest host gene TSS. Proportion (%) of insertions in sense or antisense orientations (solid grey line), upstream or downstream (dashed grey line) relative to host genes are shown. **E.** Genomic copy numbers of 126 *D. melanogaster* TE families in female dysgenic and non-dysgenic PGCs (expressed in read base pairs per TE base pairs divided by genomic coverage depth, log10). Error bars represent ± one standard deviation. Solid grey line represents perfect correlation, dashed lines indicate 5-fold change.

To assess the impact of potential biases introduced by the WGA procedure, we aligned reads to the reference genome and performed genome-wide coverage analysis over 10-kilobase windows. As expected, read coverage for single-cell samples presented more variability when compared to samples that were not subject to WGA prior to library preparation (Figure S2). However, when taking the 40 individual cells into consideration, coverage variability was shown to be randomly distributed across the genome, as previously reported (Deleye et al., 2017). Overall, a high proportion (35-95%) of the 10-kilobase windows (excluding chromosomes 4 and Y) displayed ≥10X read coverage in each analysed cell, with 44-97% of the genome displaying ≥5X read coverage.

To be able to discriminate new *P*-element insertions occurring in the genomes of isolated PGCs from those that were vertically transmitted, we characterized all insertions existing in the parental *Harwich* strain used to generate the sorted cells. To do so, we *de novo* assembled the genome of this strain using long-read Nanopore sequencing (∼75X coverage) and short-read Illumina sequencing (∼45X coverage) on bulk-extracted genomic DNA. Bioinformatic analysis revealed that the *Harwich* strain contained 32 full-length *P*-element insertions, in addition to 48 insertions corresponding to non-autonomous, internally deleted elements (Figure 2B; Table S1; Karess and Rubin, 1984; Laski et al., 1986; Rio and Rubin, 1988; Teixeira et al., 2017). We also characterised *P*-element insertions in terms of their zygosity within the *Harwich* strain, i.e. whether they were present in one or both alleles at a given locus. To do so, we used the TE discovery tool TEMP on Illumina-generated reads (Zhuang et al., 2014). This tool detects new TE insertions through comparisons of genome sequencing data to a reference genome. Since *P*-elements are absent from the reference *D. melanogaster* genome assembly (dm6), all insertions identified by TEMP represent *P*-element insertions present in the *Harwich* genome. Coverage frequencies calculated by TEMP, which serve as a proxy for the zygosity of each insertion, revealed that 58% of insertions were homozygous or nearly homozygous (coverage frequency ≥ 0.7), while the remaining 42% of insertions were segregating at variable allele frequencies (average coverage frequency = 0.3; Figure S3A).

To determine the *P*-element discovery rate in the single PGC genomes using TEMP, we focused on *P*-element insertions that are homozygous in the parental *Harwich* stock, as these insertions are expected to be present in every PGC. Taking genome coverage into account, homozygous insertions were detected at an average rate of 79% (Figure S3B). As expected, the detection rate was positively correlated with the frequency at which insertions were segregating in the *Harwich* parental strain (Figures S3B-D). Moreover, vertically transmitted insertions had coverage frequencies compatible with their expected heterozygote state in hybrid PGCs (Figure S3E). Overall, our analyses show that this approach is effective at detecting *P*-element insertions using DNA-sequencing data obtained from single PGCs.

Using this approach, we then characterised new transposition events, i.e. *P*-element insertions found in individual PGCs that were not present in the parental *Harwich* genome. Focusing on female PGCs, which inherit one copy of each chromosome from the *Harwich* and *‘white’* parental strains regardless of the direction of the cross, we found that dysgenic PGCs acquired between 0 and 5 new *P*-element insertions per diploid genome (2.2 average), only 1.5-fold more new insertions than non-dysgenic PGCs (1.5 average; Figure 2C; Table S2). Male dysgenic and non-dysgenic PGCs are not genetically identical due to the parent-of-origin inheritance of X and Y chromosomes. Despite this, and as observed for female PGCs, both dysgenic and non-dysgenic male PGCs acquired few new insertions per cell (3.6 and 2.0 average, respectively). The read coverage of new *P*-element insertions was consistent with a heterozygous state, suggesting that these insertions are present in one of the two alleles (Figure S4A).

Next, we examined the genome annotation associated with the new *P*-element insertions in the PGC genomes. Similar to what was observed for *P*-element insertions present in the *Harwich* background (Table S1), new *P*-element insertions were dispersed along the autosomes and chromosome X (Figure S4B). Based on chromatin state annotations of the reference genome (Filion et al., 2010), 60% of the new insertions were located in transcriptionally active euchromatic regions, while the remaining 40% were located in heterochromatic or transcriptionally repressed regions (Table S2). New insertions were mostly located within promoters, 5’ untranslated regions (UTRs) and first introns of genes, confirming previous findings (Table S2) (Spradling et al., 2011). By mapping their position relative to the transcription start sites (TSS) of host genes, we found that 75% of new insertions were within 1 kb of the closest TSS, mirroring what was found for insertions existing in the *Harwich* strain (Figure 2D). As was previously observed for transgenic *P*-element insertions (Spradling et al., 2011), both parental and new *P*-element insertions were also enriched in regions overlapping origins of replication (Tables S1 and S2).

Aside from *P*-elements, we sought to determine whether other transposon families become activated in PGCs during *P*-element hybrid dysgenesis. Copy numbers of 126 TE families present in the *D. melanogaster* genome were highly similar between dysgenic and non-dysgenic PGCs in both females and males (Figures 2E and S4C). This suggests that TE families other than the *P*-element are unlikely to increase in copy number in dysgenic PGCs or contribute to the germ cell loss phenotype, as previously suggested (Eggleston et al., 1988; Moon et al., 2018).

Overall, and contrary to predictions based on the insertional mutagenesis model of hybrid dysgenesis (Griffiths et al., 2000; Khurana et al., 2011), our analysis shows that dysgenic and non-dysgenic PGCs acquire similar, low numbers of new heterozygous *P*-element insertions mostly located within the promoters and introns of genes.

### PGCs are sensitive to DSBs at *P*-elements in a dosage-dependent manner

Our single-cell analysis revealed that dysgenic PGCs acquire few, scattered new *P*-element insertions, although, at the same developmental stage, virtually all dysgenic PGCs showed a strong accumulation of DSBs and cell cycle arrest (Figure 1). We hypothesised that this discrepancy could relate to the transposition mechanism employed by *P*-elements, which induces DSBs at both the excision and insertion steps. In this context, excision of multiple *P*-element copies, such as those that are typically present in *P*-strains, has the potential to induce high levels of DSBs (at least one per *P*-element copy; Bingham et al., 1982). Notably, all full-length and internally deleted *P*-element insertions in the *Harwich* background are flanked by intact TIRs, which can be recognized by the *P*-transposase and cleaved during the excision step (Figure 2B).

To test whether DSBs at *P*-elements present in the genome of the *Harwich* strain are sufficient to elicit PGC loss, we generated a transgenic line expressing the CRISPR associated protein 9 (Cas9) specifically in the germline in combination with a cassette expressing *P*-element TIR-specific guide RNA (Figure S5). The Cas9-TIR gRNA transgene was inserted into the ‘*white’* genetic background, which is devoid of *P*-elements. As expected by the absence of targets for the Cas9-TIR gRNA effector in this background, transgenic strains were homozygous viable, had normally developed ovaries, and were fully fertile (Figures 3A and 3C).

**Figure 3.**
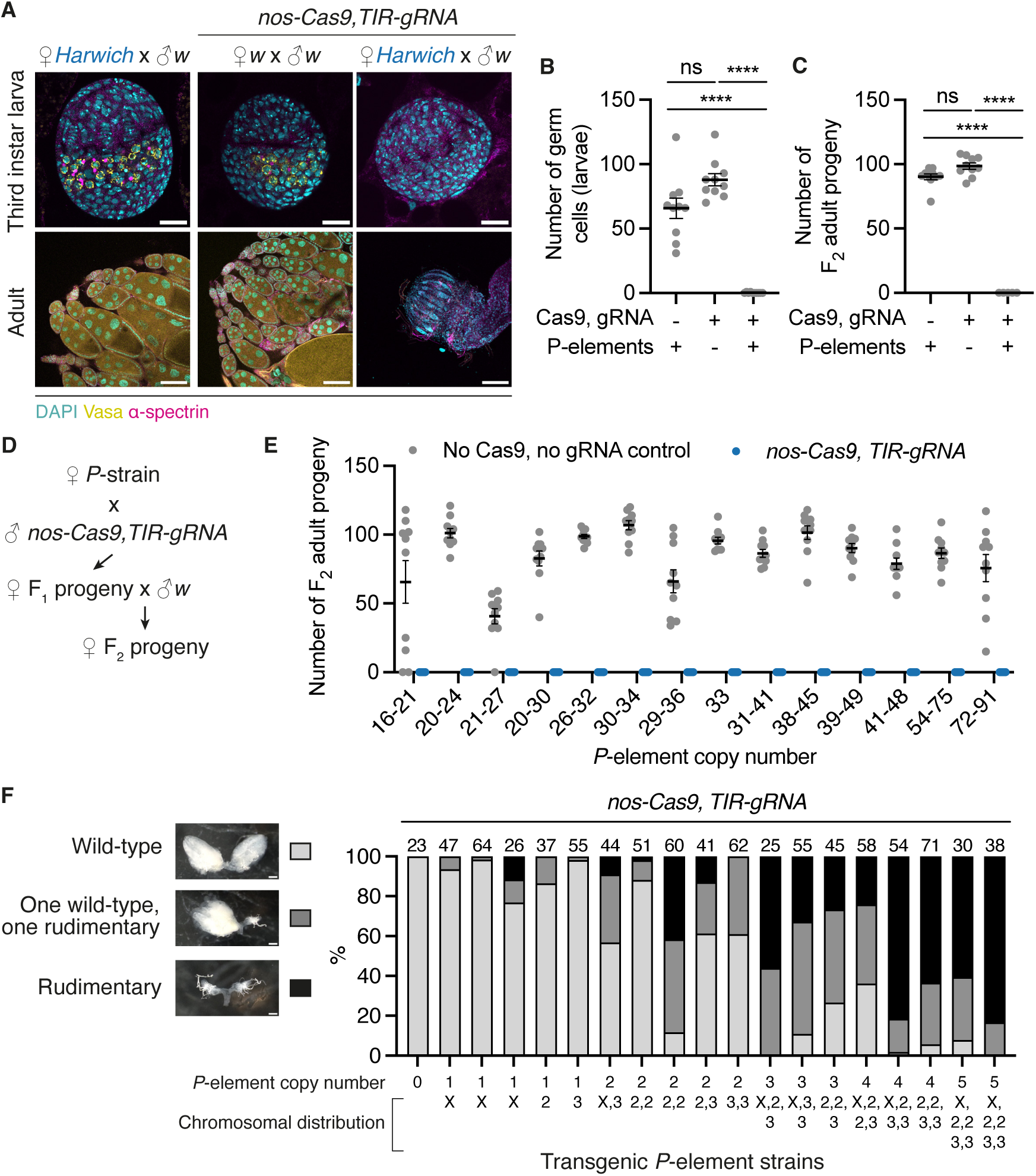
Inducing DSBs at silenced *P*-elements is sufficient to induce germ cell loss. **A.** Third instar larval ovaries (top panel) and adult ovaries (bottom panel) from non-dysgenic progeny lacking *nos-Cas9, TIR-gRNA* (left panels), progeny without *P*-elements and expressing *nos-Cas9, TIR-gRNA* (centre panels), and progeny with *P*-elements and expressing *nos-Cas9, TIR-gRNA* (right panels), labelled with DAPI (cyan), Vasa (yellow) and α-spectrin (cell borders, magenta). **B.** Number of germ cells present in larval ovaries (grey data points, n = 10) from the F1 progenies shown in (A). Black line is mean, error bars are ± SEM. **C.** Fertility of adult female progeny shown in (A) as determined by the number of F2 progeny originating from single F1 female crosses (n = 10). **** p < 0.0001, ns p > 0.05, one-way ANOVA and Tukey’s multiple comparisons test. **D.** Crossing scheme used to test if targeting Cas9 to *P*-elements in wild-derived isogenic *P*-strains, which have variable *P*-element content, modulates the penetrance of the germ cell loss phenotype observed in the *Harwich* background (A). **E.** Fertility of adult female progeny of the crosses in (D) in the presence (blue data points) or absence (grey data points) of *nos-Cas9, TIR-gRNA*, as measured by the number of F2 progeny originating from single F1 female crosses (n = 10). *P*-element copy number ranges represent estimates determined by DNA-qPCR of 5’ and 3’ regions of the *P*-element. p < 0.01 for all pairwise comparisons, unpaired *t*-test. **F.** Females from lab strains carrying between 1 and 5 transgenic *P*-element copies with varying chromosomal locations were crossed to *nos-Cas9, TIR-gRNA* males. Ovary morphology of the progeny was categorised as wild-type (light grey), one wild-type and one rudimentary ovary (dark grey) or two rudimentary ovaries (black), as shown in representative brightfield images. The number of ovary pairs analysed for each genotype is indicated above each bar. Scale bars, 20 μm (larva) or 100 μm (adult) (A), and 100 μm (F).

Targeting Cas9 to the *P*-element TIRs in non-dysgenic progeny, in which *P*-elements are present but not active, enabled us to induce DSBs in a sequence-specific manner and thereby mimic the damage caused by *P*-element excision. In contrast to what was observed in the ‘*white’* background (or in non-dysgenic control progeny lacking the Cas9-TIR gRNA transgene), when the Cas9-TIR gRNA transgene was introduced into non-dysgenic progeny, flies were viable but completely lacked germ cells and were fully sterile (Figures 3A and 3C). As we observed in hybrid dysgenesis, the germ cell loss phenotype induced by Cas9-TIR gRNA emerged during fly development, and most PGCs were lost by the third instar larval stage (Figures 3A and 3B). These results suggest that inducing DSBs at resident *P*-elements, which are present in a heterozygous state, is sufficient to elicit germ cell loss at the same full penetrance observed in hybrid dysgenesis.

Given that the *Harwich* strain contains more than 80 *P*-elements (Table S3), we asked whether a lower *P*-element copy number, and consequently a lower number of potential DSB sites, would impact the penetrance of the germline loss phenotype. Taking advantage of the Drosophila Genetic Reference Panel (DGRP) collection (Mackay et al., 2012), we selected a set of 14 wild-derived isogenic strains containing variable numbers of *P*-elements. First, we used publicly available estimates of *P*-element copy numbers that were based on analyses of sequencing data from strains in the DGRP collection (Rahman et al., 2015). We then confirmed *P*-element copy number in the subset of 14 selected DGRP stocks by quantitative PCR using primers corresponding to the 5’ and 3’ ends as well as an internal sequence only present in full-length copies (Table S3). This analysis showed that the strains contained between ∼16 and ∼91 *P*-element copies. Females from each DGRP strain were individually crossed to males carrying Cas9-TIR gRNA in the *P*-element-devoid ‘*white’* background to generate non-dysgenic progeny containing both *P*-elements and Cas9-TIR gRNA (Figure 3D). Surprisingly, non-dysgenic progeny for all these crosses were fully sterile (Figure 3E), indicating that tolerance for DSBs at heterozygous *P*-element insertions in the germline may be lower than the copy number range naturally occurring in wild strains.

Outside of their occurrence in natural strains, *P*-elements have been successfully used in the last 40 years to generate transgenic *Drosophila* lab strains (otherwise devoid of *P*-elements; Rubin and Spradling, 1982; Spradling and Rubin, 1982). For transgenesis, exogenously provided *P*-transposase is co-injected into lab strain embryos with donor vectors containing the transgenic sequences of interest, which are flanked by *P*-element TIRs (Rubin and Spradling, 1983). This process results in the random integration of TIR-flanked constructs into the host genome. Taking advantage of the extensive collections of publicly available and previously characterised transgenic strains, we used genetic crosses to combine *P*-element-derived transgenes on different chromosomes, resulting in strains carrying between 1 and 5 transgene copies and therefore much lower numbers of Cas9-TIR gRNA targets compared to wild strains (Table S3). We individually crossed females from the transgenic lab strains to *‘white’* males carrying Cas9-TIR gRNA and scored ovary morphology in the F_1_ progeny as a proxy for germline loss. Our analysis showed that the number of rudimentary ovaries, which lacked germ cells, was positively correlated with the number of transgenic *P*-elements present in the parental genome (Figure 3F). Flies carrying 1-2 transgene copies were found to be mostly fertile, while rudimentary ovaries were observed in flies carrying more than 2 transgene copies. In the presence of 5 *P*-element-derived transgenes, ovaries were mostly rudimentary, indicating that DSBs at the TIRs of as few as 5 heterozygous *P*-elements were sufficient to induce severe germ cell loss. Taken together, our results indicate that mimicking the DNA damage formed during *P*-element excision is sufficient to elicit germ cell loss at the same full penetrance observed in dysgenesis, and that the number of DSBs that PGCs tolerate is low.

### PGCs are sensitive to DSB dosage independent of *P*-elements

Given our results suggesting that PGCs tolerate few DSBs at *P*-elements, we sought to test whether this effect is a more general feature of germ cells independent of specific target sequences (*P*-element or non-*P*-element). For this purpose, we generated whole genome assemblies using long-read Nanopore (∼70X coverage) and short-read Illumina (∼50X coverage) sequencing data from a strain expressing Cas9 in the germline (*nos-Cas9*), a strain containing a docking site for gRNA-expressing transgenes (*nos-phiC31;attP2*) and a strain used to generate balanced transgenic lines (*w;TM2/TM6*). In these assemblies, we computationally searched for gRNA target sequences present in specific genomic copy numbers (Figure 4A). To avoid potential confounding effects that could arise by targeting Cas9 to coding sequences, promoters and UTRs, which may affect cell viability independently of the DSB dosage effect, we devised a tool to identify target sequences only within transposons or other non-coding sequences. We further excluded sequences present in multiple copies in close proximity to avoid inducing clustered DSBs, which could lead to rearrangements or large indels. We then performed transgenesis of these gRNA sequences (expressed ubiquitously under the control of the U6 promoter) into the strain containing a transgene docking site (*nos-phiC31;attP2*). In total, we established 12 transgenic lines expressing individual gRNAs targeting between 2 and 53 sites in the diploid genome (Table S4).

**Figure 4.**
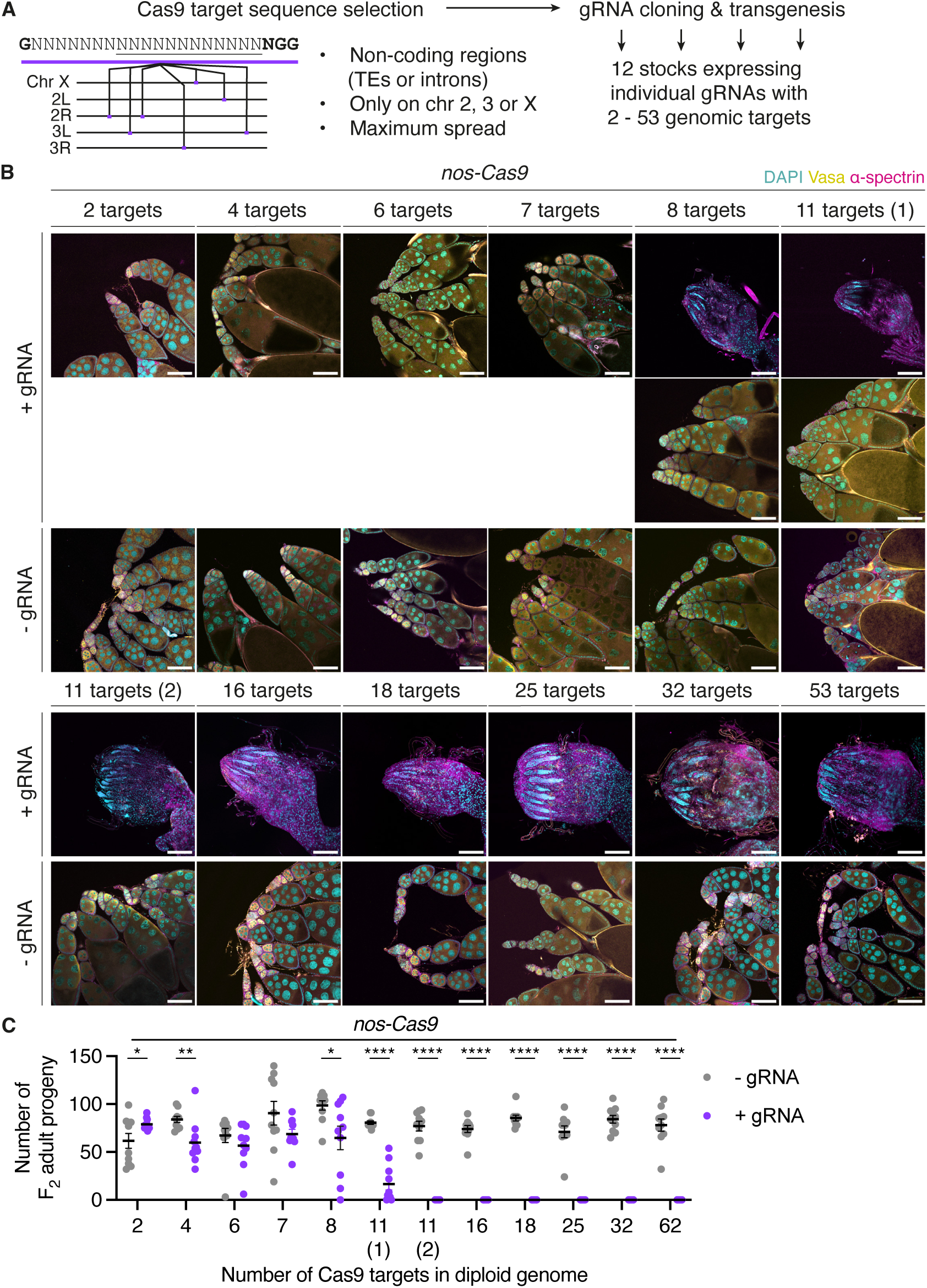
PGCs are sensitive to DSBs in a dosage-dependent manner. **A.** Cas9-based approach to systematically test DNA damage tolerance in the germline. gRNA sequences with multiple genomic copy numbers within TEs or other non-coding regions on the second, third or X-chromosome were identified in Cas9 and gRNA-expressing genomes. Individual gRNAs were separately cloned into an attB-containing vector for phiC31-mediated transgenesis, generating 12 lines expressing gRNAs with specific numbers of genomic targets in the diploid genome. **B.** Ovaries of adult F1 progeny from individual crosses of gRNA-expressing females and *nos-Cas9* males, labelled with DAPI (cyan), Vasa (yellow) and α-spectrin (magenta). Sibling progeny lacking the gRNA transgene were used as controls (-gRNA). Variable ovary morphology (wild-type and rudimentary) was observed in the presence of 8-target gRNA and one of two gRNAs with 11 targets (11-targets (1)). **C.** Fertility of the F1 progeny in (B) in the presence (purple data points) or absence (grey data points) of gRNA, as measured by the number of F2 progeny originating from single F1 female crosses (n = 10). * p ≤ 0.05, ** p ≤ 0.01, **** p ≤ 0.0001, unpaired *t*-test. Black line is mean, error bars are ± SEM. Scale bars, 100 μm.

To induce DSBs at these sites in PGCs, we individually crossed females from the gRNA-expressing lines to males from the strain expressing Cas9 in the germline (*nos-Cas9*) and assessed ovary morphology of female progeny. While ovary morphology resembled the wild-type in the presence of up to 7 Cas9 target sites in the diploid genome, increasing numbers of rudimentary ovaries were observed in females carrying gRNAs targeting 8 and 11 sites (Figure 4B). All ovaries from females carrying gRNAs targeting more than 11 sites were found to be rudimentary and completely lacked germ cells as shown by the absence of the germline marker Vasa. We assessed the effect of the germline loss on fertility and found that it was negatively correlated with the number of Cas9 target sites (Figure 4C). Targeting catalytically inactive dead Cas9 to the same sites in the germline did not affect ovary morphology, showing that Cas9-mediated DSBs, rather than its presence or binding, led to germline loss (Figure S6A). Together, these results indicate that germ cells are sensitive to DNA damage levels.

We sought to validate how efficiently the Cas9-gRNA system induced DSBs. If the system is efficient, we reasoned that targeting Cas9 to a single site in the coding sequence (CDS) of an essential gene would suffice to induce complete germ cell loss. To test this, we designed individual gRNAs targeting the CDS of three ribosomal protein (RP) genes and established transgenic stocks. Crosses between RP-gRNA and *nos-Cas9* strains produced viable female progeny with rudimentary ovaries that lacked germ cells (Figure S6B). In a small proportion of ovaries, germaria and early egg chambers were observed, but egg chambers degenerated at mid-oogenesis stages, coinciding with the nutritional checkpoint that precedes vitellogenesis (Drummond-Barbosa and Spradling, 2001). Accordingly, these flies were mostly sterile (Figure S6C). Together, our analysis shows that the Cas9 system can reproducibly induce DSBs at target sites.

To further assess the efficiency of Cas9-mediated DSB formation at the molecular level, we crossed gRNA strains with target numbers below the germ cell loss-inducing threshold (less than 11) to *nos-Cas9* males to obtain progeny with Cas9-edited germ cells. By crossing this F_1_ progeny to wild-type *‘white’* males, we obtained F_2_ flies that we expected to harbour DSB repair products at target sites. We extracted genomic DNA from 10 individual F_2_ progeny for each tested gRNA and amplified individual target sites and flanking regions by PCR. Sequencing analysis revealed that, on average, 92% of target sites were edited in the F_2_ (Figure S6D). Interestingly, in most cases, both alleles contained small indels around the target site (Figure S6E), suggesting that maternally deposited Cas9-gRNA complexes continued to target paternally inherited *‘white’* chromosomes in the F_2_ progeny, and that repair most likely occurred by non-homologous end joining (Lieber, 2010). Taken together, our molecular analysis confirms that the Cas9-gRNA system is highly efficient at inducing DSBs at its target sites.

### Mitotically dividing adult germ cells are sensitive to DSB levels

During larval development, PGCs mature into germline stem cells (GSCs; Dansereau and Lasko, 2008). In the adult ovary, GSCs continuously divide to support the production of eggs. Asymmetric GSC divisions generate differentiating daughters, which are excluded from the stem niche and thereafter undergo four mitotic cell divisions with incomplete cytokinesis to form 16-cell germline cysts (Kirilly and Xie, 2007). Meiosis begins once mitotic divisions are completed, and meiotic recombination, which involves the programmed formation and repair of DSBs, occurs at the 16-cell cyst stage while cysts are progressing through the germarium (Hughes et al., 2018). Meiotic DSBs are repaired before the 16-cell cyst exits the germarium and develops into an egg chamber containing one oocyte and 15 nurse cells, which will then further develop into a mature egg (Mehrotra and McKim, 2006). During the meiotic DSB-repair cycle, each germ cell acquires 20-24 DSBs (Mehrotra and McKim, 2006). As such, the number of meiotic DSBs acquired by adult germ cells during meiotic recombination is around double the number of Cas9 targets that we found were sufficient to result in complete germ cell loss during development (Figure 4B).

To investigate tolerance to DSBs in the adult germline prior to meiotic DSB formation, we expressed Cas9 in the *bam* domain (McKearin and Spradling, 1990; Figure 5A). The differentiation factor *bam* is expressed in a narrow developmental window during the cystoblast to 8-cell cyst stages in the germarium, when cells are still dividing mitotically and are yet to enter meiosis (Mehrotra and McKim, 2006). We used our previously characterised panel of transgenic gRNA lines targeting different numbers of genomic sites in combination with *bam-Gal4*-driven expression of *UAS-Cas9* to induce different numbers of DSBs in mitotically dividing germ cells in the adult ovary. Immunofluorescence microscopy analysis revealed that targeting up to 11 sites resulted in ovaries with wild-type morphology. However, when we targeted Cas9 to more than 11 sites, ovarioles contained germaria but had aberrant egg chamber morphology (Figure 5B). Staining for DSBs with the pH2Av antibody showed that strong DSB signal was present in cysts within the *bam* expression domain and in more posteriorly localised, further developed cysts, with the signal in these cysts likely representing meiotic DSBs (Figure 5C). We quantified the number of ovarioles containing egg chambers and found that this was negatively correlated with the number of Cas9 target sites (Figure 5D). This suggests that, like PGCs, mitotically dividing adult germ cells are sensitive to DSBs in a dosage-dependent manner, and that their tolerance threshold lies below the number of DSBs induced during meiotic recombination.

**Figure 5.**
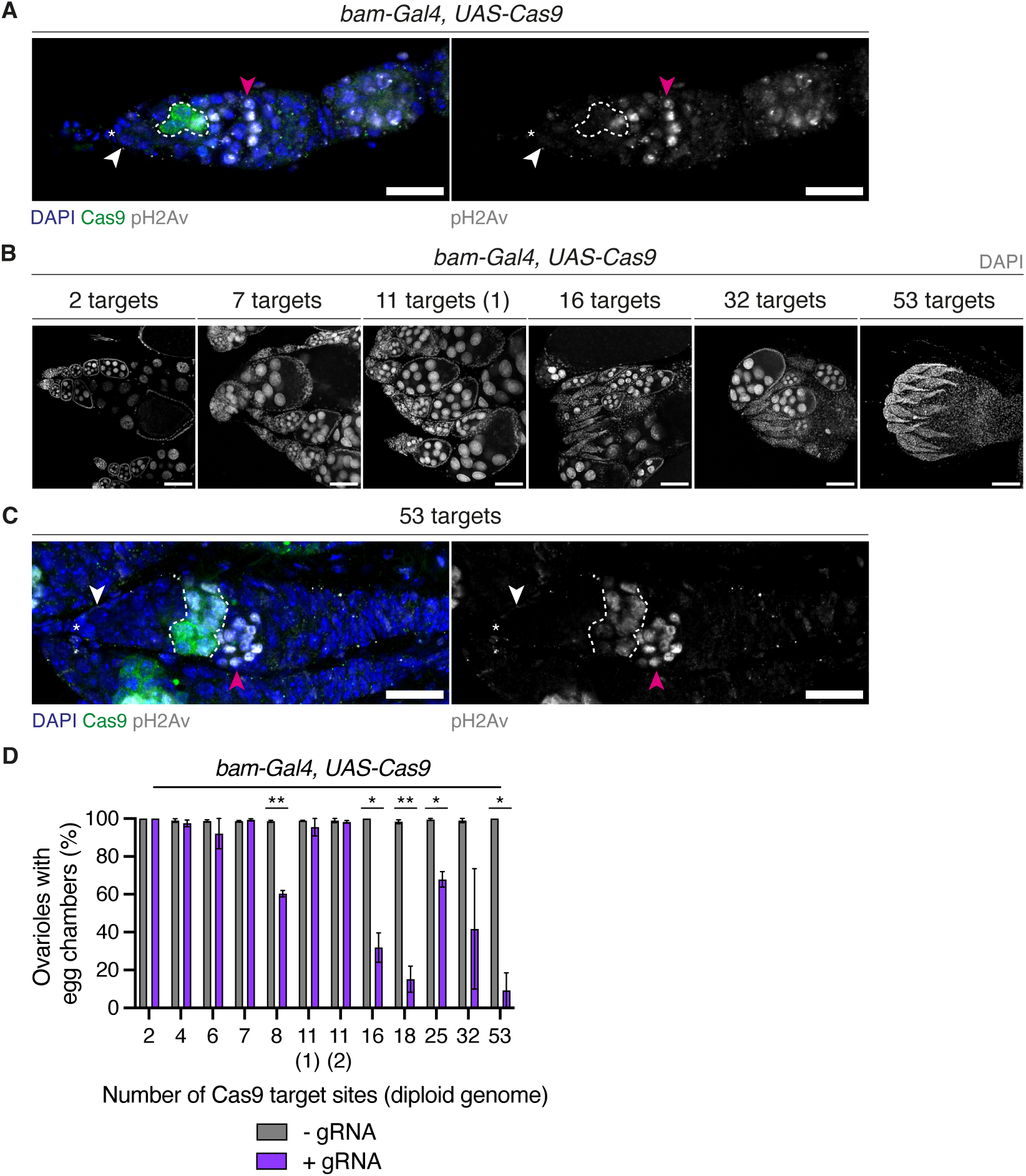
Dosage-dependent sensitivity to DSBs in pre-meiotic adult germ cells. **A.** Germarium from strain expressing Cas9 in the *bam* domain, labelled with Cas9 (green), pH2Av (grayscale and right). Asterisks indicate GSC niche, white arrowheads indicate GSCs, and magenta arrowheads indicate meiotic germ cells. Dashed line demarcates Cas9 expression domain within 2-8-cell cysts. **B.** Adult ovaries of F1 progeny from individual crosses between gRNA-expressing females and *bam-Gal4, UAS-Cas9* males, labelled with DAPI (grayscale). **C.** Germarium of F1 female in which Cas9 was targeted to 53 target sites in the *bam* domain, labelled with DAPI (blue), Cas9 (green) and pH2Av (grayscale and right). Asterisks, arrowheads and dashed lines are as in (A). **D.** Proportion of ovarioles containing egg chambers in (B), in the presence (purple bars) or absence (grey bars) of gRNA. Error bars are ± SEM. n > 150 ovarioles per genotype from two independent replicates. * p ≤ 0.05, ** p ≤ 0.01, unpaired *t*-test. Scale bars, 20 μm (A and C) or 100 μm (B).

### Post-mitotic germ cells in the adult ovary are resilient to DNA damage

To determine whether developing germ cells are equally sensitive to DSB levels after the meiotic DSB-repair cycle, we expressed Cas9 in the *TOsk* domain (ElMaghraby et al., 2022). The *TOsk-Gal4* driver is first expressed in post-mitotic germ cells (16-cell cysts) shortly after meiotic DSBs are first formed, and its expression persists throughout the remainder of oogenesis until mature eggs are formed. At the 16-cell cyst stage, the oocyte enters meiotic prophase I, where it remains until oogenesis is completed, while the nurse cells enter a specialized program of continuous, rapid cell cycles between S and G_2_ phase without cell divisions (known as endocycles), becoming polyploid to synthesize the materials required for oocyte growth (Hinnant et al., 2020; Hughes et al., 2018). To induce different amounts of DSBs in post-mitotic germ cells, we used the panel of transgenic gRNA lines targeting different numbers of genomic sites in combination with a line expressing Cas9 in the *TOsk* domain (Figure S7A). In contrast to PGCs and mitotically dividing adult germ cells, and regardless of the number of Cas9 targets (up to as many as 53), all adult female flies had morphologically wild-type ovaries that contained mature eggs (Figure S7B). We performed antibody staining against pH2Av to determine whether DSB signal was present in these ovaries. As in wild-type ovaries (Figure S7A), in both the absence and presence of gRNAs, nurse cell nuclei of early egg chambers contained pH2Av signal, which mostly disappeared from egg chamber stage 4 onwards (Figure 6A). On the other hand, and only in the presence of Cas9 and gRNAs, strong pH2Av signal was observed in oocyte nuclei of mid-stage egg chambers. Most ovarioles (62-99%) contained pH2Av-positive oocytes in the presence of gRNA targeting 6-53 sites (Figure 6B). These results suggest that, in contrast to PGCs and to mitotically dividing adult germ cells, germline development can proceed despite persistent DSBs in oocyte nuclei.

**Figure 6.**
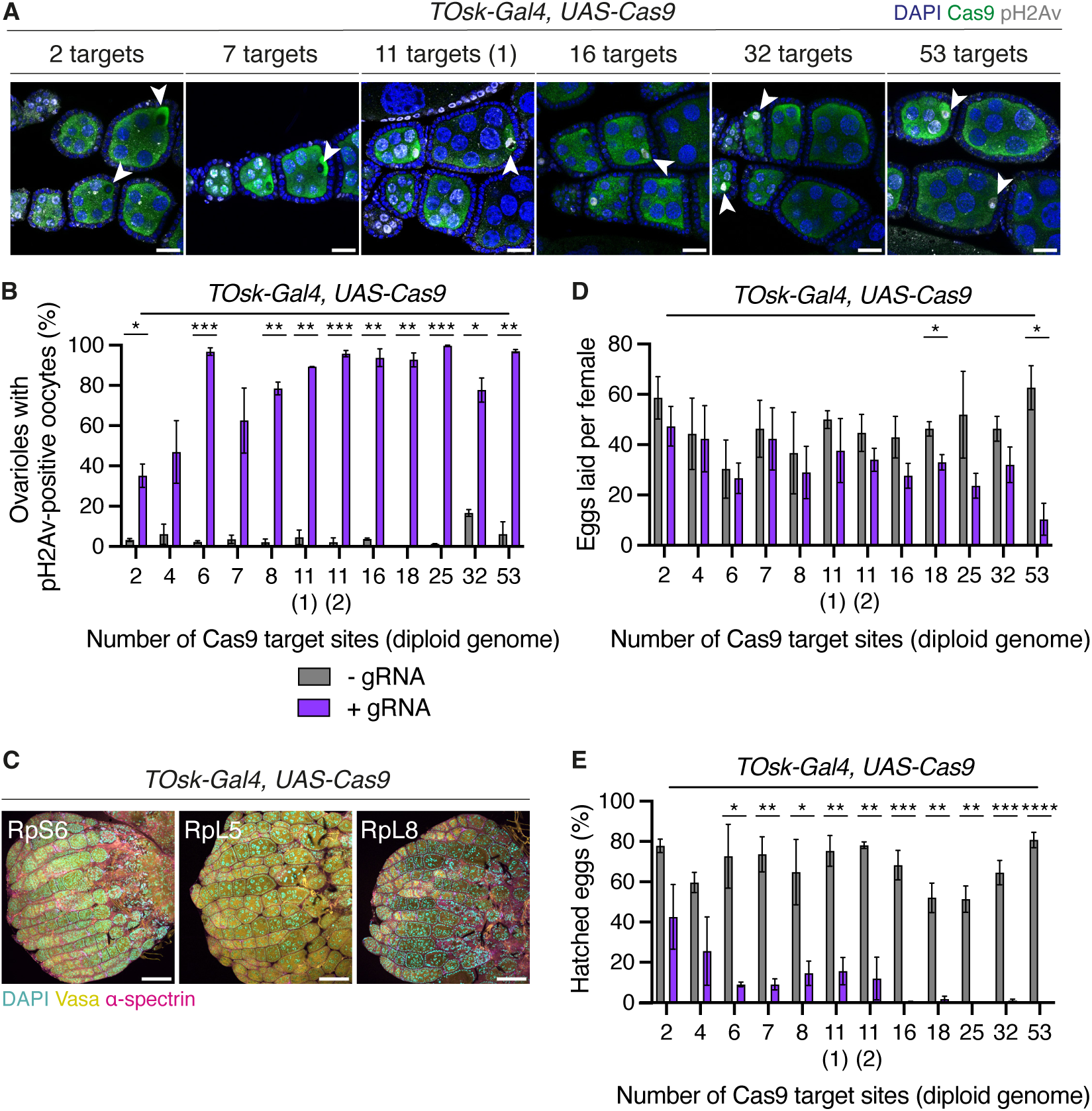
High DSB levels are tolerated following the mitotic-to-meiotic transition. **A.** Mid-stage egg chambers of F1 progeny from individual crosses between gRNA-expressing females and *TOsk-Gal4, UAS-Cas9* males, labelled with DAPI (blue), Cas9 (green) and pH2Av (grayscale). White arrowheads indicate oocytes. **B.** Proportion of ovarioles with pH2Av-positive oocytes in mid-stage egg chambers shown in (A), in the presence (purple bars) or absence (grey bars) of gRNA. n ≥ 90 ovarioles per genotype from two independent replicates. **C.** Adult ovaries of F1 progeny from individual crosses between RP-gRNA-expressing females and males expressing Cas9 in the *TOsk* domain, labelled with DAPI (cyan), Vasa (yellow) and α-spectrin (magenta). **D.** Average number of eggs laid per F1 female (n = 10, in three independent replicates) from the crosses in (A). **E.** Proportion of eggs laid by F1 females in (C) that hatched. Error bars are ± SEM. * p ≤ 0.05, ** p ≤ 0.01, *** p ≤ 0.001, **** p < 0.0001, unpaired *t*-test. Scale bars, 100 μm.

To confirm that the wild-type ovary morphology we observed was not due to the strength of expression of the *TOsk-Gal4* driver, we used our previously established RP-gRNA lines to target Cas9 to a single site within the CDS of essential genes, and assessed whether this induces germ cell loss in the *TOsk* domain. Microscopy analyses revealed that while ovaries contained wild-type germaria and early egg chambers, mid-oogenesis egg chambers had an aberrant, degenerated morphology and ovaries did not contain any mature eggs (Figure 6C). The high penetrance of these morphological aberrations indicates that the *TOsk-Gal4*-driven expression of Cas9 was sufficient to induce efficient targeting in developing egg chambers, including the polyploid nurse cell nuclei.

Regardless of the number of non-coding sites targeted by Cas9 in post-mitotic germ cells, the resulting ovaries contained eggs. To determine egg laying and hatching rates, we crossed the F_1_ progeny expressing *TOsk-Gal4* and *UAS-Cas9* (with or without gRNAs) to wild-type *‘white’* males. While the number of eggs laid in the presence or absence of gRNAs was comparable, hatching rates were reduced by 34-80% in the presence of gRNAs relative to controls (Figures 6D and 6E). We allowed eggs to develop for 16 hours, at which point PGCs have normally coalesced into embryonic gonads, and performed microscopy analysis on aged embryos. In the presence of gRNAs targeting fewer than 11 sites, embryos reached PGC coalescence stages and had wild-type morphology (Figure S7C). However, in the presence of gRNAs targeting more than 11 sites, embryos showed abnormal morphology and PGCs were mislocalised and did not coalesce to form gonads. This indicates that post-mitotic germ cells tolerate high levels of DSBs even though this may impact the development of the subsequent generation.

### DNA damage tolerance varies between mitotic and post-mitotic somatic domains

Due to its role in genetic inheritance, the germline is hypothesised to be less tolerant to DNA damage than somatic cells, favouring apoptosis over error-prone DNA repair (Bloom et al. 2019). To examine the sensitivity of somatic cells to DSB dosage, we expressed Cas9 in mitotic domains in the wing (*vg-Gal4*) and eye (*ey-Gal4*), and combined this with a subset of our transgenic lines constitutively expressing gRNAs. The *vg* phenotype, which is characterized by reduced wing size and “notched” outer wing margins (Simmonds et al., 1997), was observed in the presence of gRNAs and became more pronounced with increasing numbers of Cas9 targets (Figure 7A). However, this phenotype was consistently less severe than what we observed when driving the proapoptotic gene *rpr* in the same domain, even when targeting up to 53 sites at once. This indicates that Cas9-induced DSBs did not result in domain-wide cell death.

**Figure 7.**
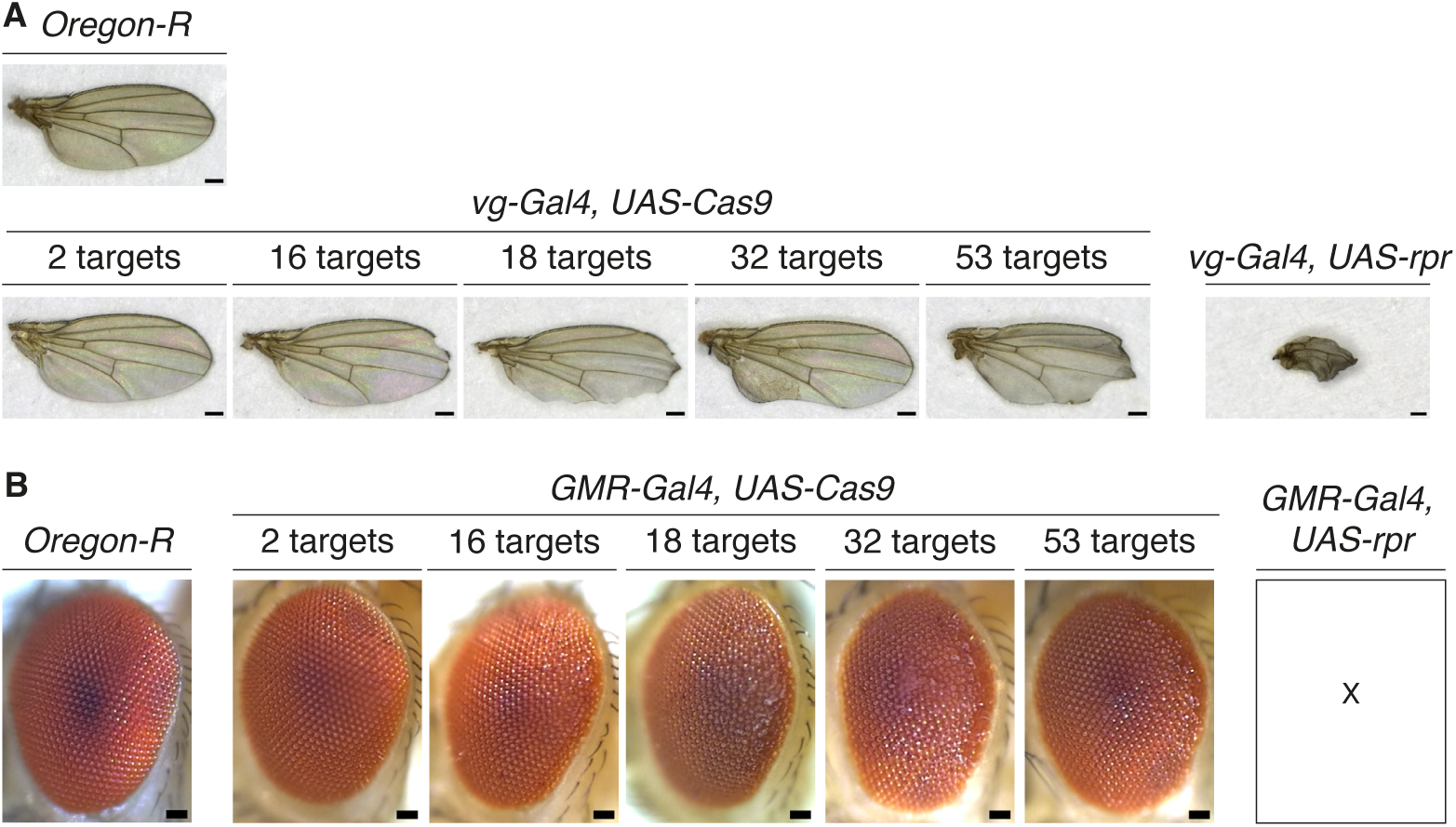
Varying tolerance to DSBs in somatic cellular domains in the wing and eye. **A.** Wing morphology of flies in which Cas9 was targeted to different numbers of genomic sites (lower left panels), or in which the apoptotic activator *rpr* was expressed (lower right panel) in the *vg* domain. Wild-type wing morphology is shown (*Oregon-R* strain, upper left panel). **B.** Eye morphology of flies in which Cas9 was targeted to different numbers of genomic sites or in which the apoptotic activator *rpr* was expressed in the *GMR* domain. X indicates no larvae, pupae or adult progeny was produced. Scale bars, 100 μm (A) and 50 μm (B).

To examine tolerance to DSBs in post-mitotic somatic cells, we expressed Cas9 in the *GMR* domain. This domain comprises cells posterior to the morphogenetic furrow that is first established in larval eye discs, and mostly contains cells in a post-mitotic or differentiating state (Freeman, 1996; Hay et al., 1994). Targeting different numbers of loci using a subset of gRNA lines in the GMR domain did not result in a marked reduction in eye size or shape regardless of the number of targets (Figure 7B). On the other hand, activating apoptosis by expressing *rpr* in the same cellular domain resulted in unviable progeny. These results suggest that post-mitotic cells in the eye may be more resilient to DNA damage compared to mitotic cells in the same tissue. Altogether, our findings indicate that responses of different cell types to DNA damage depend on the cellular differentiation state and associated cell division programs, as previously suggested (Baonza et al., 2022; Fan and Bergmann, 2014).

## Discussion

TEs successfully proliferate within eukaryotic genomes even though their activity can inflict structural and functional damage on the genome (Rahman et al., 2015; Zhang et al., 2020). Their proliferative success is conditional on their ability to mobilize within the germline. However, compromising the integrity of the germline genome threatens germ cell viability and fertility (Malone et al., 2015). Here, we used *P*-*M* hybrid dysgenesis as a model to study the effects of TE activation in the germline. Against our expectations, single-cell DNA sequencing analysis revealed that during dysgenesis, arrested embryonic PGCs only contain a few new *P*-element insertions, despite the accumulation of DSBs. Since a single source of *P*-transposase is sufficient to mobilize multiple *P*-elements in *trans* (Engels et al., 1990; O’Hare and Rubin, 1983), the activation of a single full-length *P*-element copy during dysgenesis can in principle mediate the excision of all genomic *P*-elements, including non-autonomous, internally deleted copies. Consequently, DSBs formed during *P*-element excision represent a dominant event, independent of the fact that *P*-elements are only present in the copy derived from the paternal genome during hybrid dysgenesis. Any disruption caused by random insertions would be secondary to the dominant effect of DSBs caused by excisions, as previously hypothesized (Eanes et al., 1988; Eggleston et al., 1988). This is strongly supported by our finding that mimicking DSBs formed during *P*-element excision in the absence of insertions using an engineered Cas9 system is sufficient to trigger germ cell loss during development.

Interestingly, we also observed new insertions in non-dysgenic PGCs, where *P*-element activity is largely silenced by the piRNA pathway (Aravin et al., 2007; Brennecke et al., 2008; Teixeira et al., 2017). This observation is in line with lower but nonetheless important levels of *P*-element activity, which relates to the fact that the *P*-element landscape, and that of many other TE families, is highly dynamic in natural *D. melanogaster* strains (Kofler et al., 2012; Mackay et al., 2012; Rahman et al., 2015). In this context, we propose that DNA damage tolerance thresholds may represent an important mechanism used by germ cells to control TE propagation, acting as a “last line of defence”. Our finding that DSBs at as few as 5 heterozygous transgenic *P*-elements can elicit nearly complete germ cell loss supports the idea that germ cells are unlikely to tolerate damage from more than a few *P*-elements transposing at a given time. Cells that experience any greater numbers of damage-inducing transposition events are expected to be under strong negative selection.

A further intriguing question is whether DNA damage tolerance of germ cells influences the rate of copy number expansion during *P*-element invasions into naïve strains, which initially lack piRNAs cognate to the invading TE (Bergman et al., 2017; Kelleher et al., 2018; Srivastav and Kelleher, 2017; Wang et al., 2023). The severe impact of *P*-element activity on PGC viability and exclusion of sterile individuals from selection likely present a significant hurdle during invasions. Studies of *P*-element invasions in *D. simulans* showed that populations gain ∼0.75 insertions per generation (per haploid genome) but eventually reach a copy number plateau of ∼15 insertions, at which point piRNAs cognate to the *P*-element are abundant (Kofler et al., 2022; Kofler et al., 2018). As such, the rate of *P*-element insertions during the initial stages of an invasion seems similar to the rate of new insertions we observed in dysgenic PGCs, where piRNAs are lacking. Given our results, it would be expected that during the early stages of a TE invasion, when transposition rates and DSB levels are comparatively low, *P*-element activity would be tolerated. However, as *P*-element copy numbers increase within a population, the high sensitivity of PGCs to DSB dosage would lead to an increased frequency of germ cell loss. We speculate that the germline DSB tolerance threshold, on the one hand, underlies the plateauing in copy number observed in laboratory invasions (Kofler et al., 2022; Kofler et al., 2018), and on the other hand creates the selective pressure for the TE silencing machinery against the *P*-element to become activated throughout the population. Related to this, during the hybridisation of populations with high *P*-element copy numbers and those without any *P*-elements, invading elements would only be tolerated when they are introduced *via* the maternal side, together with the protective small RNAs. In the context of DSB tolerance, the genomic *P*-element content may therefore have a major influence on the dynamics of invasions and the TE silencing machinery.

Why PGCs are so sensitive to DSBs remains unclear. DSBs are known to be particularly harmful to cells due to their propensity to cause genomic rearrangements. Moreover, their repair can introduce mutations regardless of which repair pathway is used (Chapman et al., 2012; Krenning et al., 2019). We found that PGCs are sensitive to the DSB levels, regardless of whether DSBs are induced at *P*-elements, which are largely located in promoters and introns of genes, or in other intergenic, non-coding regions. One possible explanation for this is that the DNA damage response machinery may not yet be fully matured in PGCs during the early stages of embryonic development. Alternatively, the fact that PGCs are in a state of cell cycle arrest for most of embryogenesis may prevent timely activation of the DNA damage checkpoint and recruitment of the appropriate repair machinery (Su et al., 1998). At early larval stages, when PGCs normally re-enter the cell cycle at the G_2_ to M-phase transition, the extent of genome damage may exceed the checkpoint’s capacity for repair, and despite initially prolonging cell cycle arrest, most cells are lost soon after during the first instar larval stage. The importance of the checkpoint in PGCs’ response to DNA damage is supported by the finding that *Chk2* mutants suppress some of the germ cell loss in dysgenesis (Moon et al., 2018; Xu et al., 2001). In the nematode *Caenorhabditis elegans*, PGCs trigger checkpoint-induced apoptosis in response to persistent UV-induced DNA lesions (Ou et al., 2019), suggesting that this checkpoint-mediated response is conserved. Of note, in *Drosophila* larvae, mutations affecting DNA repair and the checkpoint, or repair alone, result in hypersensitivity to irradiation-induced DNA damage, but mutations affecting only the checkpoint do not (Jaklevic and Su, 2004). Therefore, there are likely other, cell cycle-interacting factors that facilitate DNA damage sensing and repair in PGCs. Transcriptome analysis of PGCs may help to obtain an unbiased and holistic view of the molecular pathways that are activated during dysgenesis.

A further open question is whether PGCs are sensitive to different sources of DNA damage. DNA cleavage by *P*-transposase is known to generate staggered-ended breaks with 17-bp overhangs, while Cas9 induces staggered-ended DNA breaks with 1-3-bp overhangs (Beall and Rio, 1997; Shou et al., 2018; Zuo and Liu, 2016). It is possible that different types of breaks and repair outcomes may trigger different cellular responses. Our molecular analysis of DSB repair products at Cas9 target sites showed that Cas9-induced DSBs in PGCs were most likely repaired by non-homologous end-joining. By contrast, *P*-element excision sites are frequently repaired by homologous recombination (Engels et al., 1990). Previous work has shown that the choice of repair pathway after *P*-element excision depends on the developmental stage (Preston et al., 2006). Whether repair outcomes and cellular responses to DSBs more generally depend on when damage is induced during development remains to be studied.

In contrast to PGCs and mitotically dividing adult germline cysts, germ cells showed remarkable resilience to DSBs at the post-meiotic stage, where DSB repair is essential to resolve the obligatory meiotic crossovers. When DSBs were induced at this stage, we found that ovary morphology and egg laying were unaffected even though oocytes contained strong DSB signal, indicating that oogenesis can be completed despite high levels of genome damage. We hypothesise that this may be related to the fact that germ cells have already been exposed to meiotic DSBs and repair by this stage of oogenesis. Meiotic DSB repair during early-to-mid-prophase I involves a specific set of factors (Hughes et al., 2018), but when the repair process fails, cells trigger a meiotic checkpoint that involves the canonical ATM pathway factors *mei-41* and *Chk2* (Abdu et al., 2002; Ghabrial et al., 1998). We did not observe the axial patterning defects associated with meiotic checkpoint activation (Abdu et al., 2002; Shim et al., 2014), indicating that DSBs induced after the meiotic DSB-repair process may not trigger this checkpoint. Therefore, and despite the fact that the DSBs we induced resulted in aberrant zygotic development, our findings are in line with a much lower sensitivity or even an absence of checkpoint-related DNA damage responses in the post-mitotic germline.

With respect to TEs, which must mobilize in germ cells to increase in copy number in the heritable genetic material, the comparatively greater tolerance of post-mitotic germ cells to genome damage may have implications for proliferation strategies. Despite high levels of DNA damage endured once meiotic DSBs are repaired, oogenesis can be completed and, on rare occasions, embryos develop and hatch in the subsequent generation. The post-mitotic domain may therefore provide the most suitable developmental window for TEs to become active and thereby increase their chances of propagating across generations. Importantly, it has been shown that retrotransposon transcripts produced in the nurse cells can be shuttled to the transcriptionally silenced oocyte *via* microtubules (Van De Bor et al., 2005; Wang et al., 2018). As such, it is possible that TEs use the nurse cell genome as a platform for transposition in the post-mitotic germline.

Finally, another open question is whether the shifts in tolerance to DSBs between different stages of germline development is a specific feature of this cell lineage, or rather a phenomenon that occurs in other cell lineages during fate specification and differentiation. We observed lower tolerance to equivalent DNA damage levels in mitotic primordial domains in the wing, as compared to post-mitotic domains within the eye. One explanation for this observation is that the primordial cells may not yet be equipped to cope with genome damage to the same extent as fully differentiated cells. Alternatively, their cell cycle state and associated checkpoints may be driving the decision of whether to repair or die.

## Experimental Methods

### *Drosophila* genetics and husbandry

*D. melanogaster* stocks were maintained on standard cornmeal medium at 18°C. Flies used for genetic crosses were kept on propionic medium supplemented with yeast at 25°C. For genetic crosses, virgin females and males were added to fresh vials and allowed to lay for 2-3 days at 25°C. To induce dysgenesis, crosses were established at 29°C (Kidwell et al., 1977). To assess F_1_ female ovary morphology, adult female progeny of the appropriate genotype were collected and kept on fresh vials supplemented with yeast for 1-2 days to “fatten” ovaries prior to dissection. To assess the ability of F_1_ females to generate progeny, 2-6-days-old F_1_ females were collected, individually crossed to two *w^1118^* males and allowed to lay for 2-3 days. The number of F_2_ progeny (male and female) emerging from 10 individual crosses was assessed 12 days after crosses were set up (Teixeira et al., 2017).

For egg laying and hatching assays, 10 adult females of the tested phenotype were mated with 5 ‘wild-type’ *w^1118^* males in fresh vials for 2 days before being transferred to standard embryo collection cages for a further 2 days with apple juice/agar plates changed every 24 hours. The total number of eggs laid per cage over the next 24 hours was counted, and agar plates containing eggs were incubated at 25°C for a further 24 hours prior to the total number of hatched eggs being determined. Egg-laying and hatching assays were performed in three biological replicates for each genotype and corresponding control.

*D. melanogaster* stocks used were:

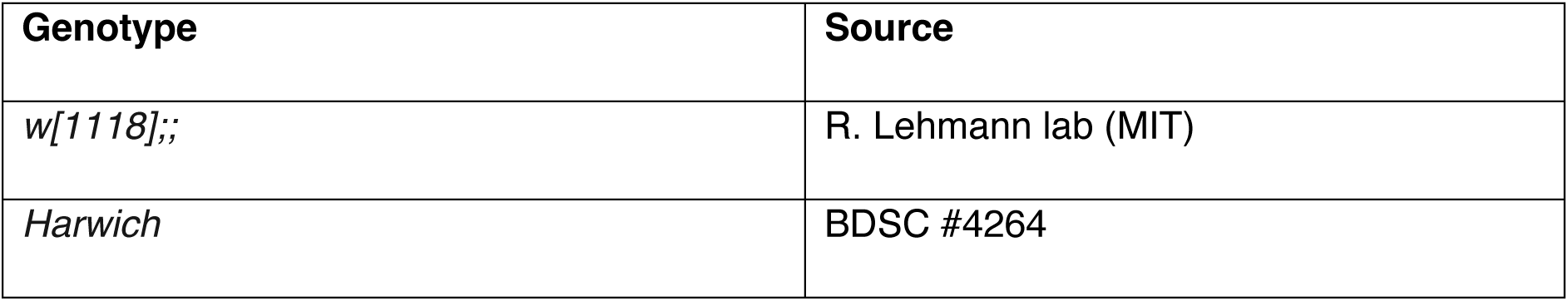
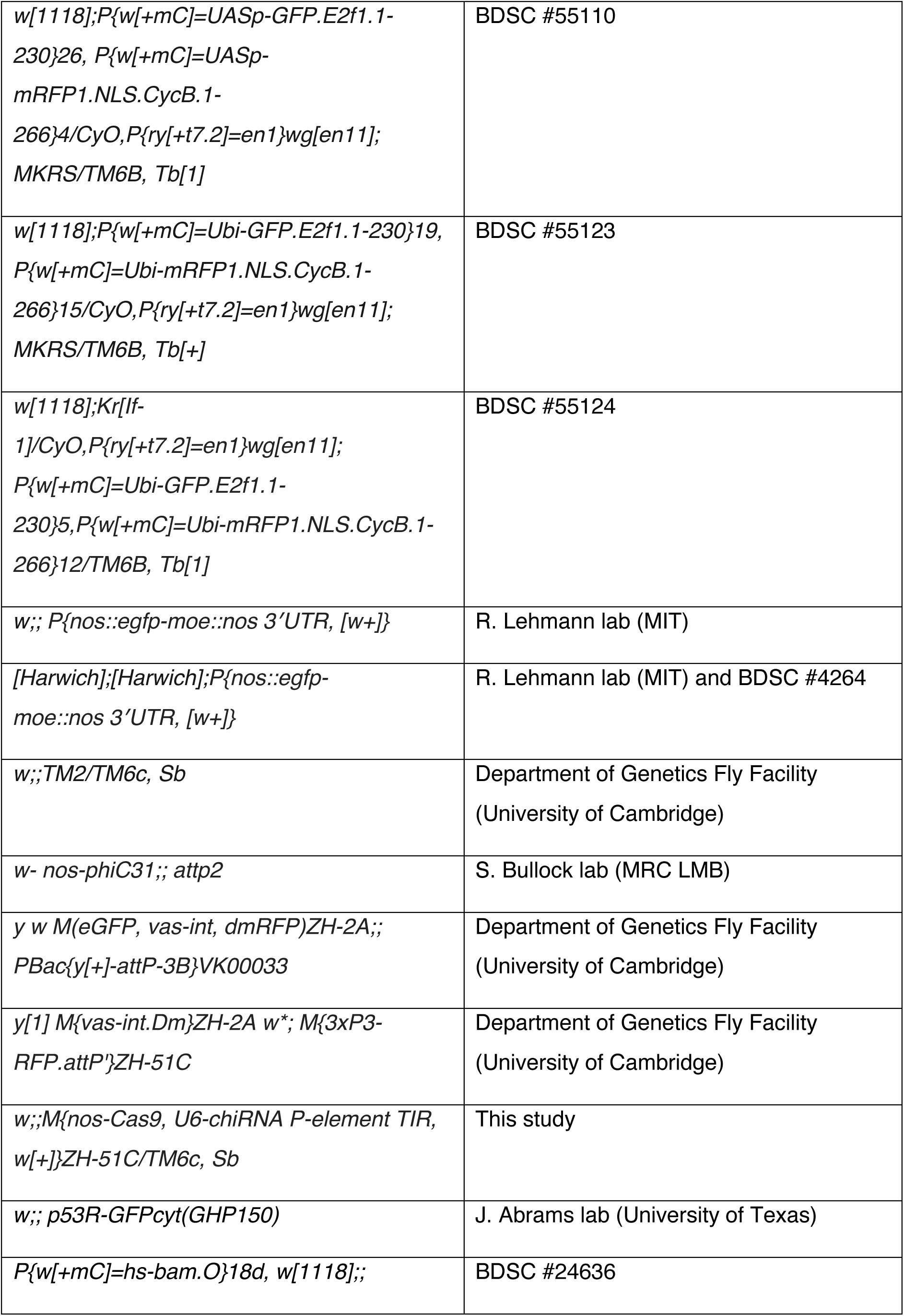
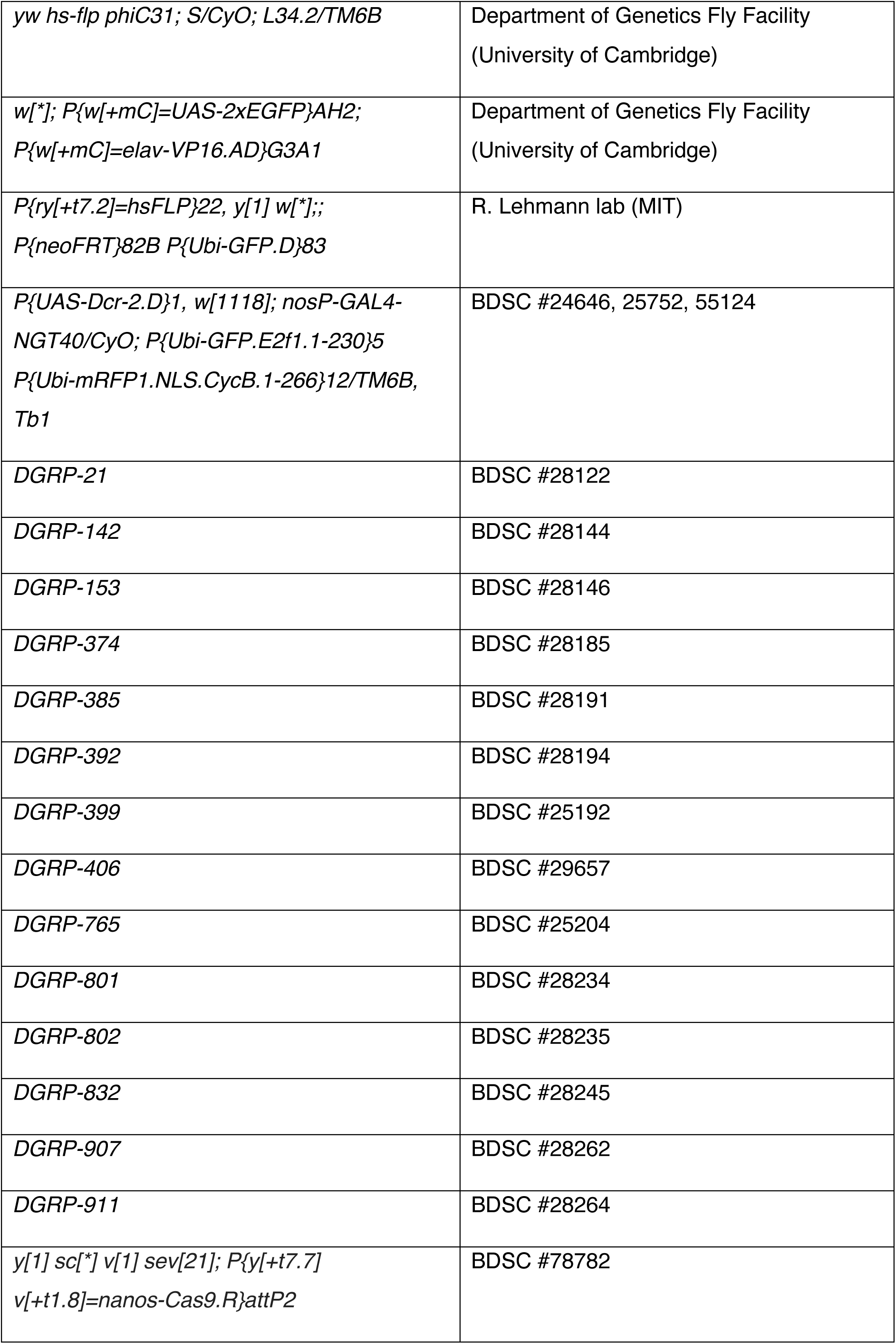
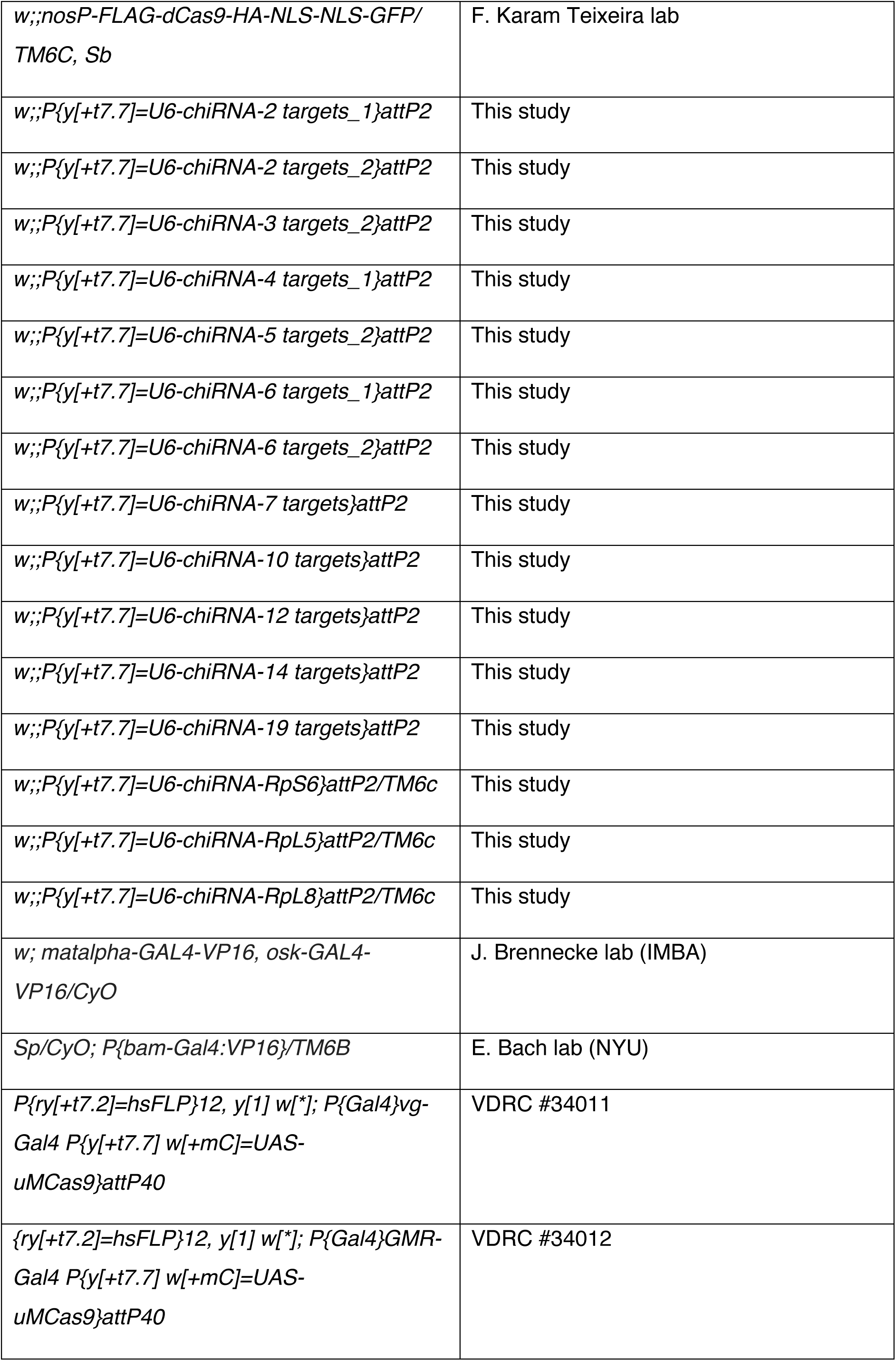
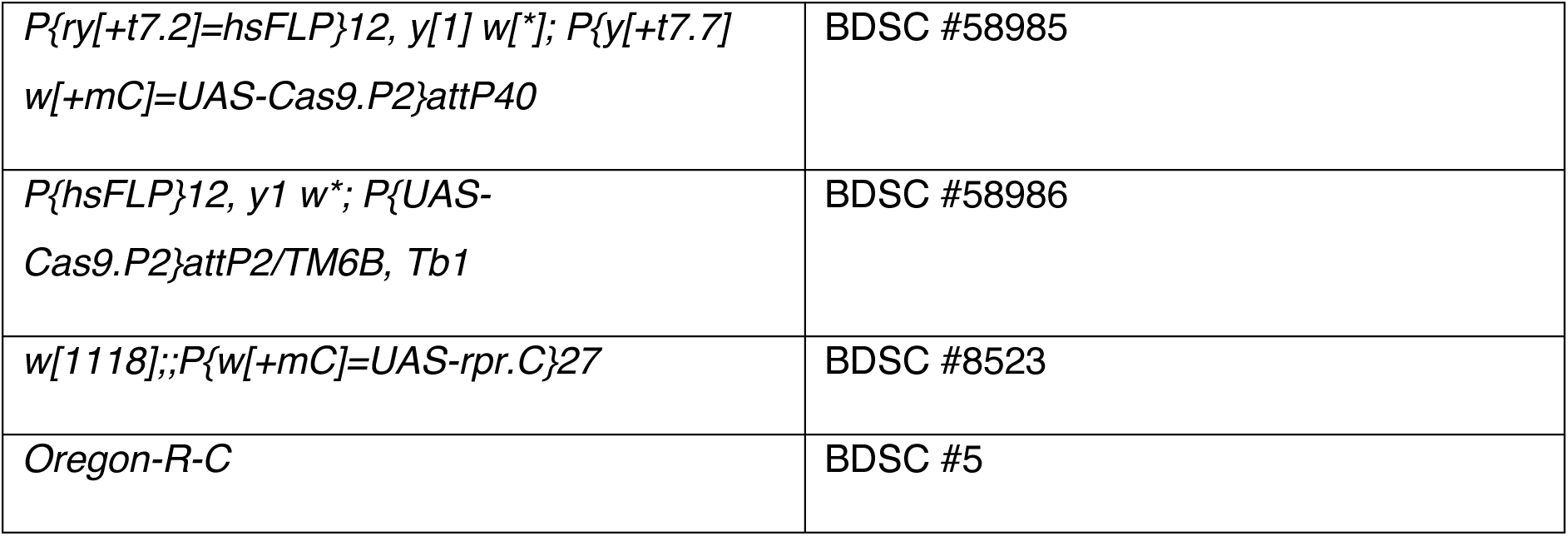

### Immunofluorescence and confocal microscopy

0-14h hour old embryos were collected on standard apple agar plates supplemented with yeast paste. Embryos were dechorionated in 2.5% bleach for two minutes and subsequently incubated in fixation solution containing 9% formaldehyde in 1X PBS and heptane (1:5) for 30 minutes (Seifert and Lehmann, 2012). Fixed embryos were hand-devitellinized in PBT (1X PBS with 0.1% Triton X-100) and blocked in PBTB (1X PBS with 0.2% Triton X-100 and 1% Bovine Serum Albumin) for 1 hour. Larvae were collected at 24-48h (first instar) and 72-120h (third instar) and dissected in ice cold 1X PBS (Teixeira et al., 2017). Larval tissue was fixed in 4% formaldehyde for 20 minutes, washed three times in PBT (1X PBS with 1% Triton X-100) and blocked in PBTB for 1 hour (Maimon and Gilboa, 2011). Adult ovaries were dissected in cold 1X PBS and fixed in 4% formaldehyde for 20 minutes. Ovaries were washed three times in PBTB for 20 minutes each and blocked in PBTB for 1 hour. Blocked samples were incubated in primary antibodies diluted in PBTB overnight at 4°C. Samples were washed in PBTB three times, then incubated in secondary antibodies diluted in PBTB for 2 hours at room temperature in the dark. Samples were washed in PBTB three times and mounted in Vectashield medium containing DAPI (Vector Labs). Fluorescent images were acquired on a Leica SP8 confocal microscope using 20X dry or 40X oil objectives.

To quantitate fluorescent signal, Z-stacked images of embryonic and larval gonads were acquired. Female embryonic gonads were identified by the presence of PGCs and by absence of Vasa-positive male-specific somatic gonadal precursor cells (Renault, 2012). To quantify fluorescent signal in embryonic and larval PGCs, images were loaded into Fiji ImageJ (Schindelin et al., 2012). Boundaries of PGC and epidermal cell nuclei were defined as regions of interest (ROIs). For each ROI, area, mean intensity, and area integrated density were measured on GFP and RFP channels. To account for background signal, measurements were also taken on non-epidermal somatic cells (RFP- and GFP-negative) within the same image slices. For each PGC and epidermal cell ROI, the corrected total cell fluorescence (CTCF) was calculated as previously described (McCloy et al., 2014).

A Leica epifluorescence microscope fitted with an LED light source and 488 nm filter was used to quantify pH2Av-positive oocytes. The number of ovarioles containing at least one stage 2-6 egg chamber with pH2Av signal was counted manually across two biological replicates.

| **Antibody** | **Concentration** | **Source** | **Research Resource Identifier (RRID)** |
| --- | --- | --- | --- |
| Rabbit anti-Vasa | 1:5000 | R. Lehmann lab (MIT) | AB_2940894 |
| Rat anti-Vasa | 1:50 | DSHB | AB_760351 |
| Mouse anti-pH2Av | 1:200 | DSHB (UNC93-5.2.1-s) | AB_2618077 |
| Rat anti-RFP | 1:500 | Chromotek (RMA5F8) | AB_2336064 |
| Chicken anti-GFP | 1:500 | Aves Labs (GFP-1010) | AB_2307313 |
| Mouse anti-alpha-spectrin | 1:200 | DSHB (3A9 323/M10-2) | AB_528473 |
| Rabbit anti-Cas9 | 1:1000 | Diagenode (C15310258-100) | AB_2715516 |
| Alexa Fluor Plus 488 goat anti-mouse | 1:250 | Invitrogen (A32723) | AB_2633275 |
| Alexa Fluor 647-AffiniPure donkey anti-rabbit | 1:250 | Strattech (711-605-152-JIR) | AB_2492288 |
| Alexa Fluor 488 goat anti-chicken | 1:250 | Invitrogen (A11039) | AB_2534096 |
| Alexa Fluor 594 donkey anti-rat | 1:250 | Invitrogen (A21209) | AB_2535795 |
| Cy3-AffiniPure donkey anti-mouse | 1:250 | Strattech (715-165-150-JIR) | AB_2340813 |

### Isolation of PGCs from embryos

Reciprocal dysgenic and non-dysgenic crosses between the *Harwich* and *w^1118^* strains expressing the fluorescent PGC marker transgene (*P(nos::egfp-moe::nos-3’UTR[w+])*) were established in separate cages (∼250 flies per cage) fitted with apple agar plates supplemented with yeast paste at 25°C for 2 days (Sano et al., 2005; Teixeira et al., 2017). In parallel, cages of the *w^1118^* stock (lacking the germ cell marker) were established as a negative control. Cages were moved to 29°C to induce dysgenesis for 1 day prior to collection. Embryos were collected for 4 hours and aged at 29°C. 11–16-hour old embryos were dechorionated in 2.5% bleach for 2 minutes, washed with water and incubated in cell sorting buffer (“Balanced Saline”) for 2 minutes (Chan and Gehring, 1971). Embryos were manually dissociated by Dounce homogenization in 5 mL cold cell sorting buffer. The homogenate was sequentially filtered through 100 μm and 20 μm cell strainers (Celltrics). Live/dead staining with Zombie Aqua fluorescent dye (Biolegend) was used to exclude dead cells. The dye was reconstituted in DMSO according to the manufacturer’s instructions and diluted 1:100 in PBS. 1 mL of cell homogenate was transferred to separate microcentrifuge tubes as controls for live/dead staining. Cells were pelleted, washed, and resuspended in cold 1X PBS (unstained controls) or dye solution and incubated at room temperature for 10 minutes. Cells were washed twice and resuspended in cold cell sorting buffer. Cells were sorted on a BD FACS Aria Fusion cytometer at the NIHR Cambridge Biomedical Research Centre Cell Phenotyping Hub facility. 488 nm and 568 nm lasers were used to identify the GFP-positive, Zombie Aqua-negative cell population. Single live, GFP-positive cells were sorted into REPLI-g cell storage buffer (Qiagen).

### DNA extraction and amplification

Genomic DNA from freshly sorted single cells was extracted and amplified using the REPLI-g Advanced Single Cell kit (Qiagen) according to the manufacturer’s instructions. Whole genome-amplified DNA was quantified using the Qubit dsDNA Broad Range Assay kit (Invitrogen). Size distribution of amplicons was determined using the TapeStation Genomic DNA Screen Tape assay (Agilent). Bulk genomic DNA was extracted from 10 non-virgin female adult flies using the Quick-DNA Microprep kit (Zymo) and quantified by Qubit. Bulk-extracted DNA was not amplified prior to library preparation.

### Quantitative PCR screen to determine cell sex

To distinguish male (XY) and female (XX) sorted PGCs, a quantitative PCR (qPCR) screen was set up based on the presence or absence of the Y chromosome. Sets of primers corresponding to the CDS of Y chromosome genes *ARY* and *FDY* were designed using Flybase and primer3. Sets of primers corresponding to genes on the autosomes (*Dmn*, *Und* and *nos*) were used as copy number controls. qPCR reactions were set up in technical duplicates using LightCycler 480 SYBR Green I Master Mix (Roche), 5 ng amplified DNA and 1 μM forward and reverse primers and run on a LightCycler 480 machine (Roche) using the following protocol: 10 minutes at 95°C; 10 seconds at 95°C, 1 minute at 60°C (45 cycles). Melting curves were used to verify primer specificity. Female cells were identified by lack or low amplification of Y-chromosome genes (high C_T_ values and unspecific priming on melting curve) as compared to autosomal genes. Oligonucleotides used were:

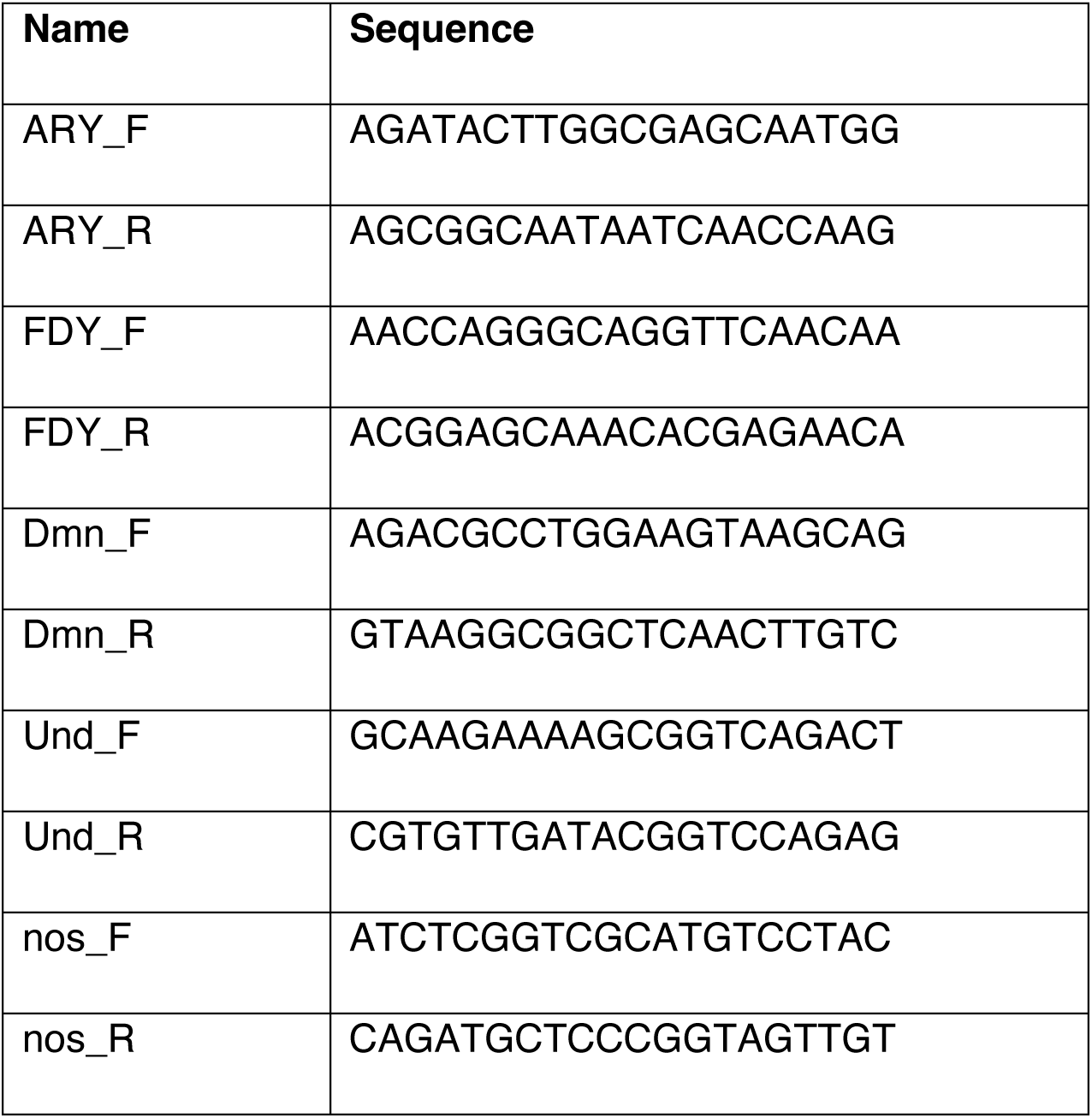

### Short-read DNA-sequencing

DNA-sequencing libraries were prepared using the DNA Prep kit (Illumina) with 500 ng of whole genome-amplified DNA (single PGC samples) or bulk DNA (whole-fly, no amplification) as input. Libraries were multiplexed using Nextera index adapter oligos (Illumina) and quantified by Qubit (Invitrogen). Library size distribution was determined by Bioanalyzer (Agilent). Multiplexed libraries were sequenced on a NovaSeq 6000 system (Illumina) as paired-end, 150-nt-long reads.

Reads were mapped to the reference genome dm6 using BWA MEM (Li and Durbin, 2009). Alignments were filtered for unique mappers, and a map quality cut-off of 30 was applied using Samtools (Li et al., 2009). The total number of mapped reads and mapped read count per chromosome were obtained using the Samtools utility idxstats. Overall genome coverage was calculated by multiplying the total mapped read count by paired-end read length (2 x 150-nt) and dividing by the dm6 reference genome size. To obtain read coverage over 10-kb windows, dm6 was indexed and sorted using Samtools and command line utility sort. Bedtools was used to define coordinates of 10-kb windows (autosomes and X-chromosome) and determine the number of reads per window for each sample (Quinlan and Hall, 2010). Coverage per window was determined by multiplying the read count per window by paired-end read length and dividing by bin size. The (k,e)-mappability of the reference genome dm6 was obtained using genmap, with a k-mer size of 150 and mismatch number of 2 (Pockrandt et al., 2020). To obtain the average mappability per 10-kb window, genome-wide mappability scores of 150 k-mers for each 10-kb window were determined using the Bedtools tool map with the option mean.

### Long-read DNA-sequencing

High Molecular Weight (HMW) genomic DNA was extracted using the Genomic Tips 100/G kit (Qiagen) following a modified version of the manufacturer’s instructions. For each genotype of interest, 60 female flies were flash-frozen in liquid nitrogen and homogenized in lysis buffer, vortexed and incubated at 37°C for 1 hour. Following incubation with Proteinase K, samples were centrifuged at 5,000xg, 4°C for 10 minutes. After the addition of isopropanol, DNA eluate was centrifuged at 10,000xg, 4°C for 30 minutes. Purified DNA was resuspended in Elution Buffer (Qiagen) and dissolved at 37°C for 1 hour and at 4°C for >12 hours. Concentration and purity (A_260_:A_280_) were determined by Qubit and Nanodrop (Thermo Fisher). DNA size range was determined using TapeStation (Agilent).

Libraries for long-read sequencing were prepared using 3-7 μg HMW DNA as input for the Oxford Nanopore Technologies (ONT) Ligation Sequencing kit (SQK-LSK110) according to the manufacturer’s instructions with the following modifications. DNA repair and end-prep reactions were incubated at 20°C for 30 minutes and 65°C for 30 minutes. Samples were diluted in 90 μl Elution Buffer (NEB), mixed with 120 μl AMPure XP beads (Beckman Coulter) and incubated on a rotator mixer at room temperature for 10 minutes. Bead-bound DNA was washed twice with 300 μl 80% ethanol. Beads were resuspended in 63 μl Elution Buffer (NEB) and incubated at 34°C for 30 minutes prior to sample elution. End-prepped DNA was quantified using the Qubit dsDNA Broad Range kit. Sequencing adapter ligation reactions were incubated at room temperature for 1 hour. Libraries were subsequently diluted in 50 μl Elution Buffer for bead clean-up as described above. Beads were resuspended in 26 μl Elution Buffer (ONT) and incubated at 34°C for 30 minutes prior to sample elution. Final libraries were quantified by Qubit.

Libraries were sequenced on SpotON Flow Cells (R9 version, FLO-MIN106D) primed using the Flow Cell Priming Kit (EXP-FLP002) on a MinION Mk1C sequencing device (ONT) according to the manufacturer’s instructions. To generate sufficient genome coverage, two genotypes were run on each flow cell for ∼24 hours each. Flow cells were washed between runs using the Flow Cell Wash kit (EXP-WSH004).

### Genome assembly

ONT reads that passed quality control (Q score ≥ 8) were used as input for the Flye genome assembler (Kolmogorov et al., 2019). ONT reads were then used to polish the Flye output draft assembly using the Medaka sequence correction tool (ONT) and scaffolded on the dm6 reference genome using the D-GENIES alignment tool (Cabanettes and Klopp, 2018). Assemblies were further polished using Illumina short-read data for three consecutive runs of the Racon polishing tool (Vaser et al., 2017).

### Transposon and *P*-element insertion analysis

To identify existing *P*-element insertions in the *Harwich* strain used for hybrid dysgenesis crosses, the search_repeat_copies Perl script was run with the *P*-element consensus sequence (Flybase) and genome assembly as inputs (Gebert et al., 2021). In parallel, TEMP was run on Illumina data from the *Harwich* strain and sorted PGCs (Zhuang et al., 2014).

Insertions identified by TEMP and supported with <2 reads were removed from analyses. To validate the presence and determine precise coordinates of *P*-element insertions relative to dm6, short-read alignments generated with BWA MEM, and long-read alignments generated with minimap2 (Li, 2018) were loaded into the Integrative Genomics Viewer IGV (Robinson et al., 2011). Soft-clipped reads overlapping the insertion site were extracted and insertion coordinates were determined by BLAST against the *P*-element consensus sequence and dm6 (Flybase). Coverage frequencies for insertions with ≥10 supporting reads were obtained from the variant support values output by TEMP.

Total genomic TE copy numbers were determined by mapping reads to the complete set of *D. melanogaster* TE consensus sequences (Flybase) using BWA MEM and Samtools with map quality ≥ 30. To normalize read counts for each TE family by genome coverage, read counts were multiplied by the paired-end read length and divided by TE consensus length and genome coverage (Gebert et al., 2021). Mean normalized TE counts in dysgenic and non-dysgenic female and male PGC genomes were determined.

### DNA qPCR for copy number validation

Genomic DNA was extracted from individual female adult flies from stocks of interest using QuickExtract solution (Lucigen) according to the manufacturer’s protocol. Primer sets for the *P*-element 5’ and 3’ TIRs and internal region were designed using the *P*-element consensus sequence (Flybase) and primer3. For qPCR reactions, 1 μL of DNA was added to a master mix containing 2X LightCycler 480 SYBR Green I Master Mix, forward and reverse primers (both 1 μM) and nuclease-free water. For each genotype and primer set, reactions were performed in 2 technical replicates. Autosomal genes, *Dmn* and *nos*, were used to benchmark copy number. qPCR reactions were run as described above. Threshold cycle (C_T_) values were averaged across replicates. For each sample, C_t_ values for the two positive control genes were averaged. For each of the three *P*-element regions, C_T_ values were subtracted from the average autosomal gene C_T_ value (Δ C_T_ autosomal gene - Δ C_T_ *P*-element). Copy number estimates of the full-length, 5’ and 3’ TIR amplicons were calculated as 2^-(Δ^ ^C^_T_ ^autosomal gene - Δ C^_T *P*-element)._

Oligonucleotides used were:

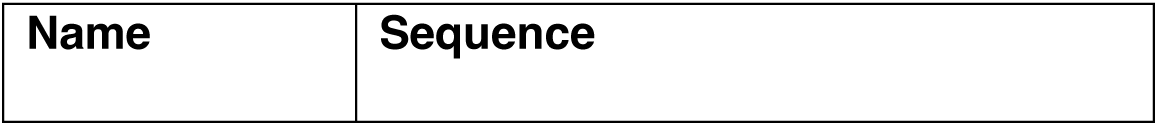
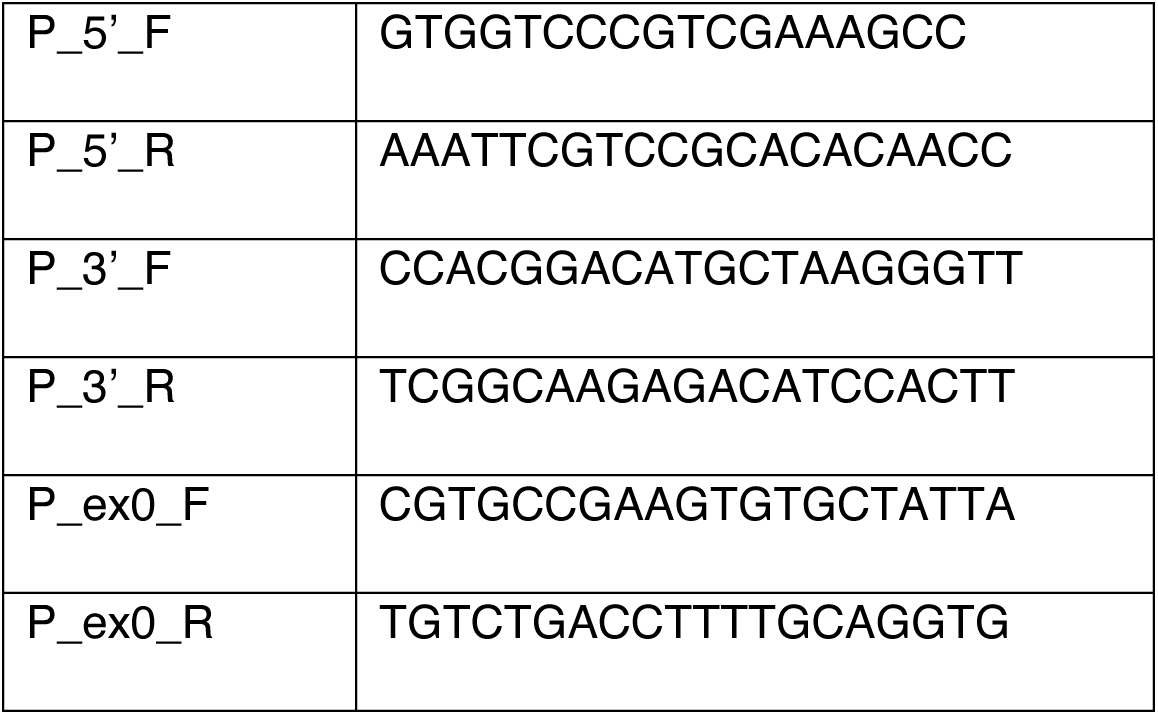

### CRISPR-Cas9 target sequence selection

A gRNA sequence containing a Protospacer Adjacent Motif (PAM) was identified within the *P*-element TIRs (1-20 bp and 2888-2907 bp of the full-length *P*-element sequence) using the *P*-element consensus sequence (Flybase) and published gRNA design guidelines (Port et al., 2014).

We established a Perl-based programme (GenoScythe) that performs a heuristic search for gRNA sequences with different numbers of genomic targets within dm6. Genomic positions for introns and TEs were extracted from GTF and RepeatMasker files with annotations based on dm6 (Flybase). An initial list of 500,000 sequences was generated by extracting 20-nt short sequences from random positions within genomic intron and TE regions containing a 5’-G and followed by a PAM according to published gRNA design guidelines (Port et al., 2014). Sequences with stretches of at least four simple repeats (e.g. AAAA or ACACACAC) were excluded. The number of sequences originating from each chromosome was proportionate to the corresponding chromosome length. Unique sequences were given identifiers and mapped to the reference genome using Bowtie (Langmead et al., 2009). Genomic hits were then counted for alignments with a perfect match in the 12 nucleotides adjacent to the PAM (“proximal” part) and with a maximum of 2 mismatches in the 8 nucleotides furthest from the PAM (“distal” part). Sequences with off-target hits, e.g. outside introns and TEs, and sequences with hits that had mismatches were filtered out at this stage. Similarly, sequences with any hits outside of the specified set of chromosomes were discarded. The final output tables contained the sequence, total number of hits, number of hits on each chromosome arm, hit coordinates and strand.

Presence of the gRNA sequences targeting multiple genomic sites was validated within the genomes of the *nos-int;P(attP2)* and *w;TM2/TM6* strains used to generate balanced gRNA-expressing lines, and the *nos-Cas9* strain. Briefly, for each strain, short-read (Illumina) and long-read (ONT) DNA-sequencing data was obtained as described above. Genomes were assembled and polished as described above. The set of dm6-derived target sequences was mapped to each genome assembly using Bowtie. The resulting SAM files contained the number of hits, location, and strand (+ or -) for each target sequence within each assembly. The number of diploid target sites was determined as double the average haploid number of target sites across the three genomes.

RP genes with an associated *Minute* (haploinsufficiency) phenotype were identified from the literature (Marygold et al., 2007). CRISPR Optimal Target Finder (Gratz et al., 2014) was used to identify Cas9 target sequences within the CDS for each gene (Flybase).

gRNA target sequences (including PAMs) were:

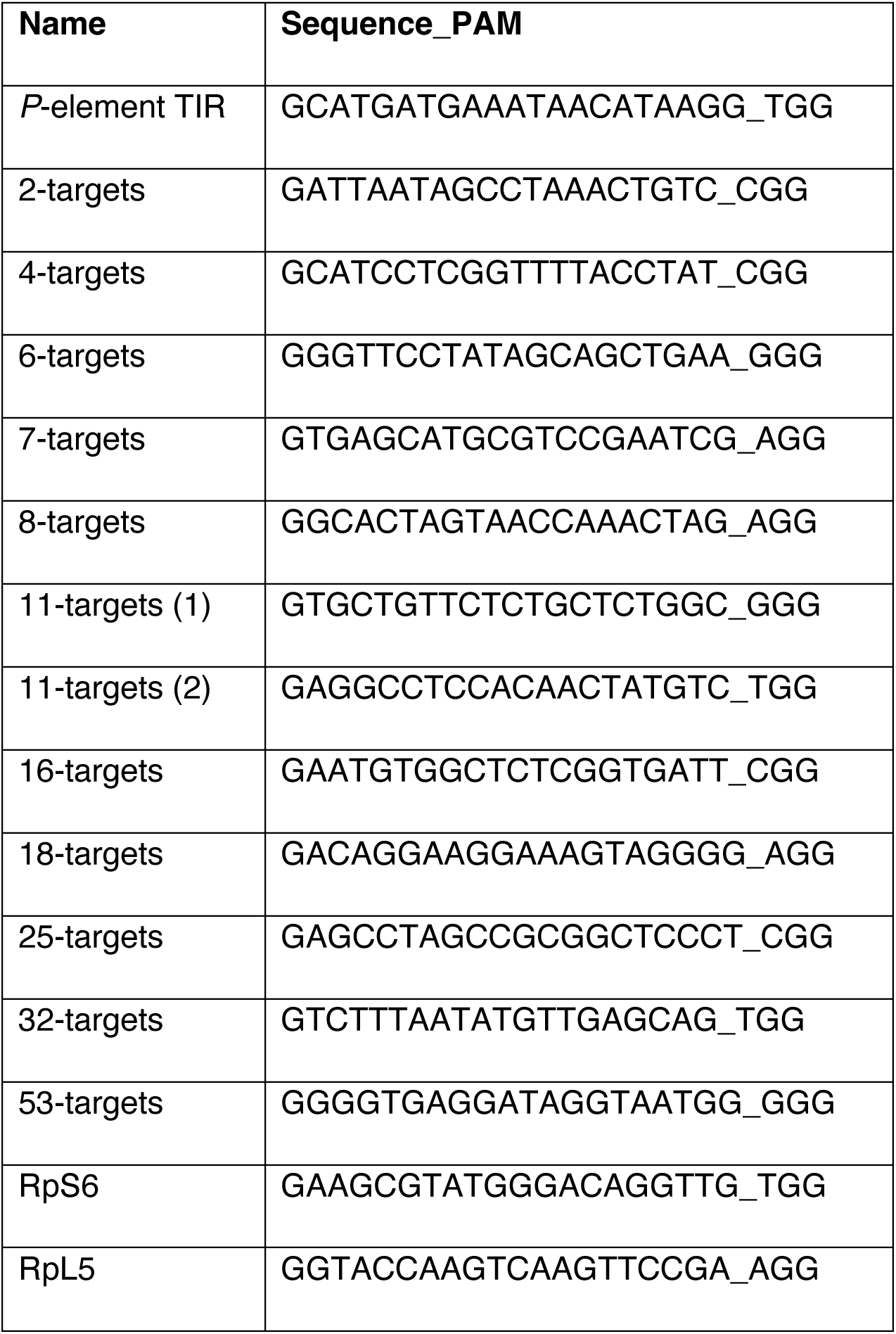
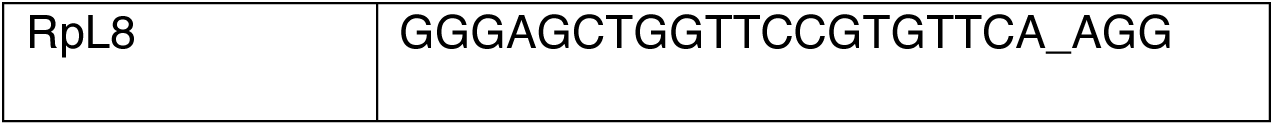

### CRISPR-Cas9 target sequence cloning and transgenesis

The *TIR-gRNA* sequence was cloned into the pU6-BbsI-chiRNA plasmid (Addgene #45946) using the FlyCRISPR protocol (Port et al., 2014). The resulting pU6-chiRNA cassette was amplified by PCR and inserted into the pnos-Cas9-nos plasmid (Addgene #62208) at the NheI restriction site. gRNA sequences targeting different numbers of genomic sites and RP-gRNA sequences were individually cloned into the pU6-BbsI-chiRNA plasmid. gRNA cassettes were amplified by PCR and individually inserted into a pWalium22 backbone containing a *mini-white* selection marker and attB site (Drosophila Genomics Resource Center #1473). Final constructs were assembled by Gibson assembly (NEB) according to the manufacturer’s protocol. Plasmid assembly was confirmed by Sanger sequencing and restriction digestion analysis. Plasmid DNA was injected into embryos from the *nos-int;PBac(attP9-A)* stock (for transgenesis of the TIR-gRNA, nos-Cas9 construct), and from the *nos-int;P(attP2)* stock (for transgenesis of the multi-target gRNA and RP-gRNA constructs) at the Department of Genetics Fly Facility. Final, balanced transgenic lines were generated by backcrosses to *w;TM2/TM6* flies.

### CRISPR-Cas9 molecular validation

For molecular analysis of DSB repair products at Cas9 target sites, PCR primers were designed to amplify ∼250-2,200 bp regions containing the target sequences and flanking regions for “hit” locations present in all of the *nos-Cas9*, *nos-int;attP2*, and *w;TM2/TM6* genome assemblies (Primer3). To validate PCR primer specificity and efficiency, bulk-extracted genomic DNA from these strains was used to set up PCR reactions for each primer set using Quick Load Taq 2X Master Mix (NEB) according to the manufacturer’s protocol. Amplicon size was validated by gel electrophoresis. Amplified DNA was purified using the QIAquick PCR Purification kit (Qiagen) prior to Sanger sequencing analysis. To evaluate Cas9 editing efficiency, F_1_ virgin females from crosses between the *nos-Cas9* line and gRNA lines with 2-11 target sites (below the full germ cell loss inducing threshold) were crossed to *w^1118^* males. F_2_ adult females were collected for DNA extraction using QuickExtract solution, and PCR amplification and Sanger sequencing analysis were performed as described above on ≥10 biological replicates per target site. Sequences perfectly matching the wild-type strains were classified as unedited, while sequences containing indels and mutations around the Cas9 target site were defined as edited.

Oligonucleotides used for validation were:

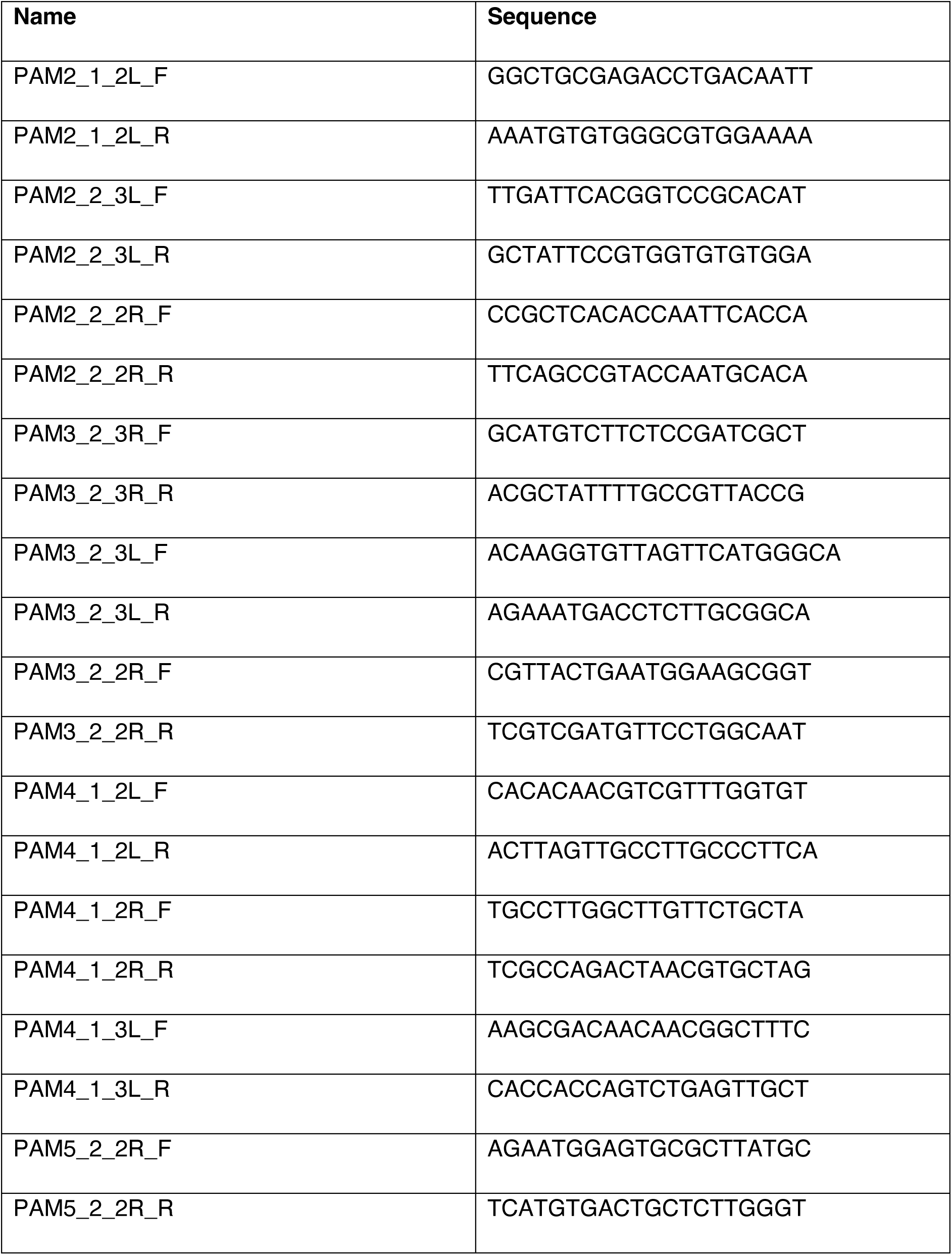
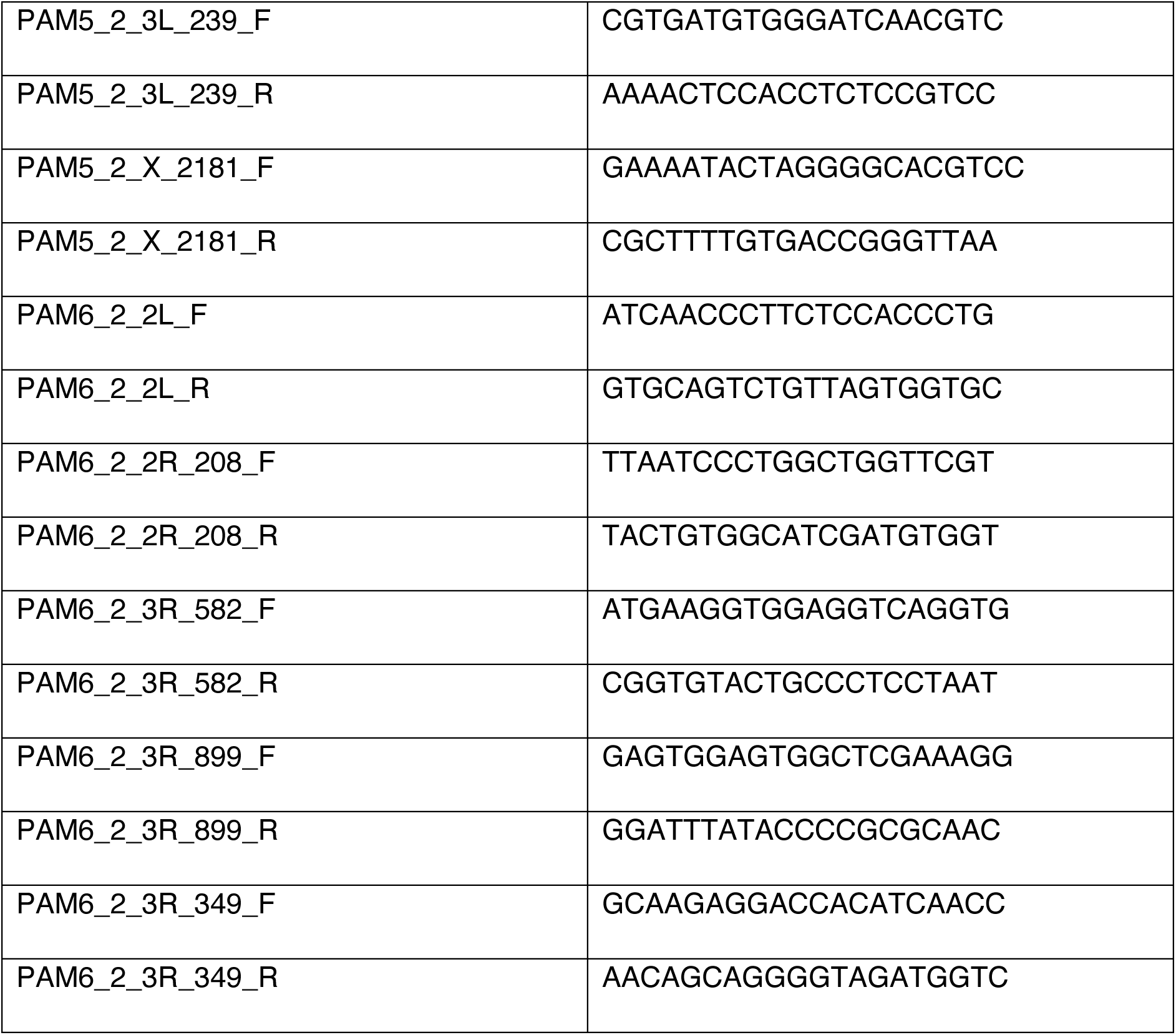

### Statistical analysis

All experiments were conducted at least three independent times. Statistical analysis was performed using GraphPad Prism software. Statistical significance was tested by unpaired *t*-tests. Pearson correlation coefficient was determined with 95% confidence intervals. No statistical methods were used to predetermine sample size. Experiments were neither intentionally randomized nor intentionally ordered. Investigators were not blinded to allocation during experiments and outcome assessment.

## Resource Availability

### Lead contact

Further information and requests for resources or reagents should be directed to the lead contact, Felipe Karam Teixeira.

### Materials availability

Transgenic *Drosophila* lines generated in this study are available upon request.

### Data and code availability

DNA-sequencing data and genome assemblies generated in this study is deposited at the NCBI Sequence Read Archive (SRA) (https://www.ncbi.nlm.nih.gov/) under project PRJNA1044074. Custom code used in this study is deposited at Github (https://github.com/d-gebert/GenoScythe). Any additional information required to reanalyse the data reported in this study is available upon request.

## Acknowledgements

We thank R. Lehmann and R. Durbin for antibodies and reagents; J. Abrams, E. Bach, J. Brennecke, F. Jiggins, R. Lehmann, H. Ma, C. O’Kane, the Department of Genetics Fly Facility, the Vienna *Drosophila* Resource Center and the Bloomington *Drosophila* Stock Center for fly reagents; the Cambridge NIHR BRC Cell Phenotyping Hub and the Department of Genetics imaging facility for technical support; B. Fischer for technical assistance; and S. Russell and P. Andersen for discussions. This work was supported by the Wellcome Trust and Royal Society Sir Henry Dale Fellowship 206257/Z/17/Z (to F.K.T.), Human Frontier Science Program CDA-00032/2018 (to F.K.T.), Wellcome Trust Studentship 220026/Z/19/Z (to G.J.), Walter Benjamin Postdoctoral Fellowship from the Deutsche Forschungsgemeinschaft (to D.G.), Public Service Department of Malaysia Scholarship (to T.R.K.) and and a Biotechnology and Biological Sciences Research Council (BBSRC) Doctoral Training Partnership (to E.S.). For the purpose of Open Access, the author has applied a CC BY public copyright license to any Author Accepted Manuscript (AAM) version arising from this submission.

## Author contributions

G.J. and F.K.T. designed the experiments. G.J., D.G. and F.K.T. performed bioinformatic analyses. G.J., T.R.K., E.S. and S.M. performed all further experimental analyses. G.J. and F.K.T. wrote the manuscript.

## Supplemental Information

**Figure S1.**
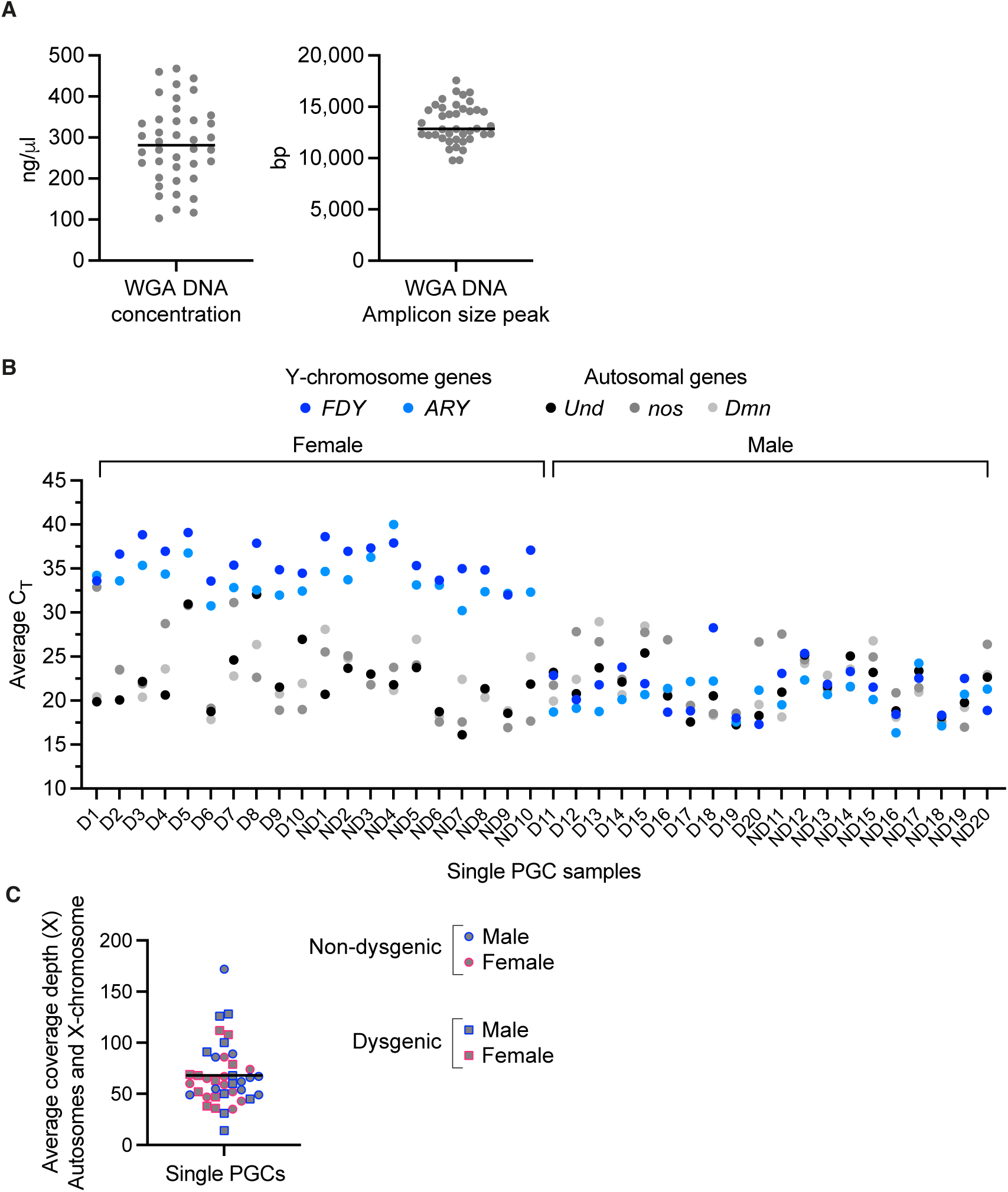
WGA DNA from single sorted PGCs generates high overall genome coverage in DNA-sequencing data and can be used to determine cell sex. **A.** Concentration (left panel) and amplicon size peak (right panel) of WGA DNA from sorted PGCs. Each data point represents one PGC sample. Black lines indicate mean. B. Average threshold amplification cycle (CT) values for Y-chromosome genes *ARY* (light blue) and *FDY* (dark blue) and autosomal genes *Dmn* (light grey), *nos* (dark grey) and *Und* (black) for the 40 PGC samples, as determined by qPCR on WGA DNA. High CT values (≥30) for Y-chromosome genes designate female PGCs, low CT values (<30) designate male PGCs. C. Average coverage depth across the autosomes and X-chromosome in 40 whole-genome PGC data sets. Dysgenic (squares) and non-dysgenic (circles), male (blue) and female (magenta) PGCs are indicated. Black line represents mean.

**Figure S2.**
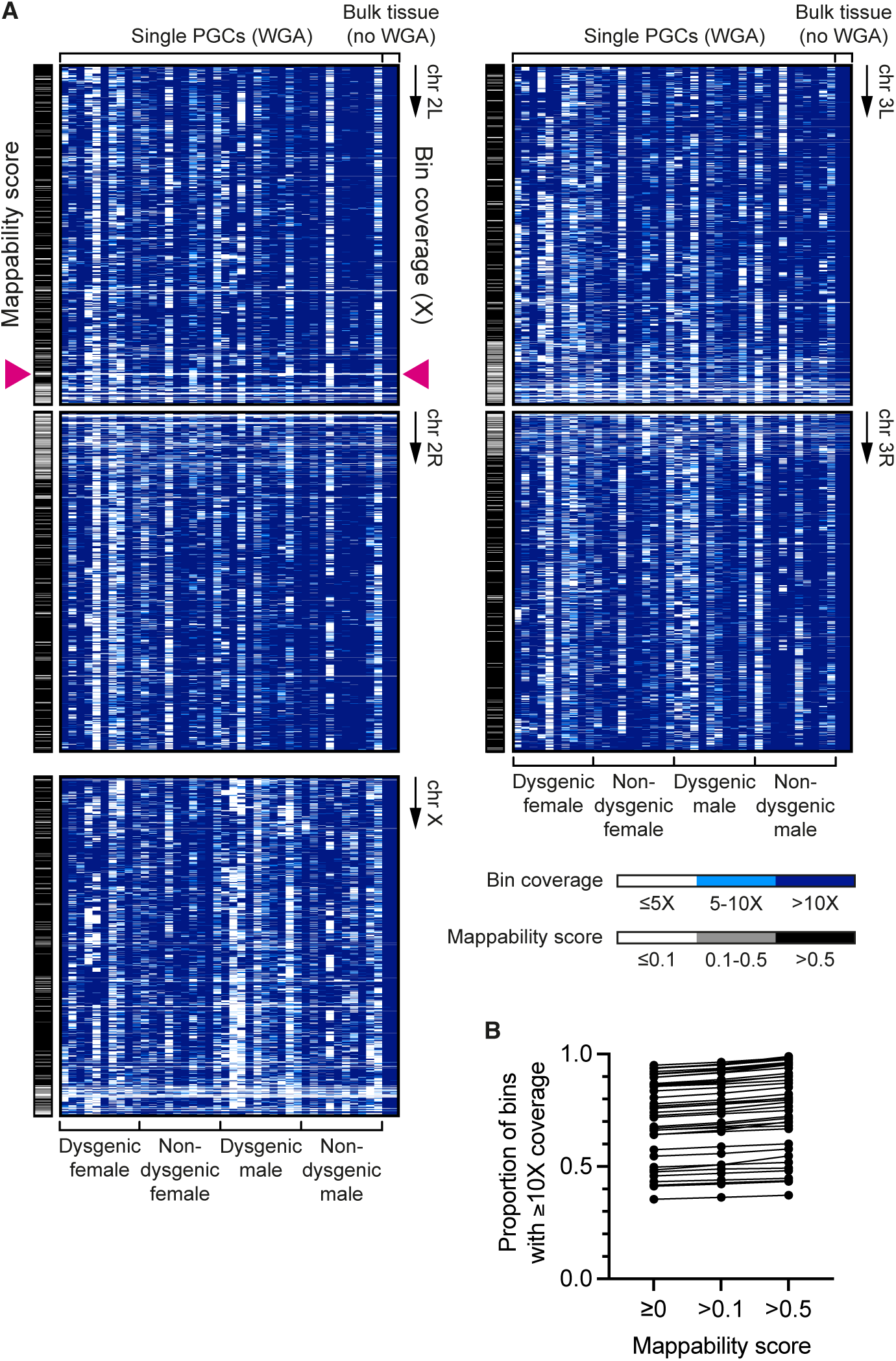
Whole-genome sequencing of WGA DNA from single PGCs produces variable genome coverage. **A.** Heatmaps showing coverage depth (white, <5X; light blue, <10X; ≥10X, dark blue) over 10-kb genomic regions along chromosome arms 2L, 2R, 3L and 3R and the X-chromosome (rows) for 40 PGCs and 2 non-WGA (bulk tissue) samples (columns). Mappability scores (white, <0.1; grey, 0.1-0.5; black, >0.5) for 10-kb windows based on the *dm6* reference genome are shown to the left of each coverage heatmap. Magenta arrowheads indicate a region with low mappability with low coverage depth in all samples, as one of few exceptions to the overall random distribution of coverage variability in PGC data sets. **B.** Proportion of 10-kb windows with ≥10X read coverage when 3 different minimum mappability scores were applied. Data points represent individual PGC genomes.

**Figure S3.**
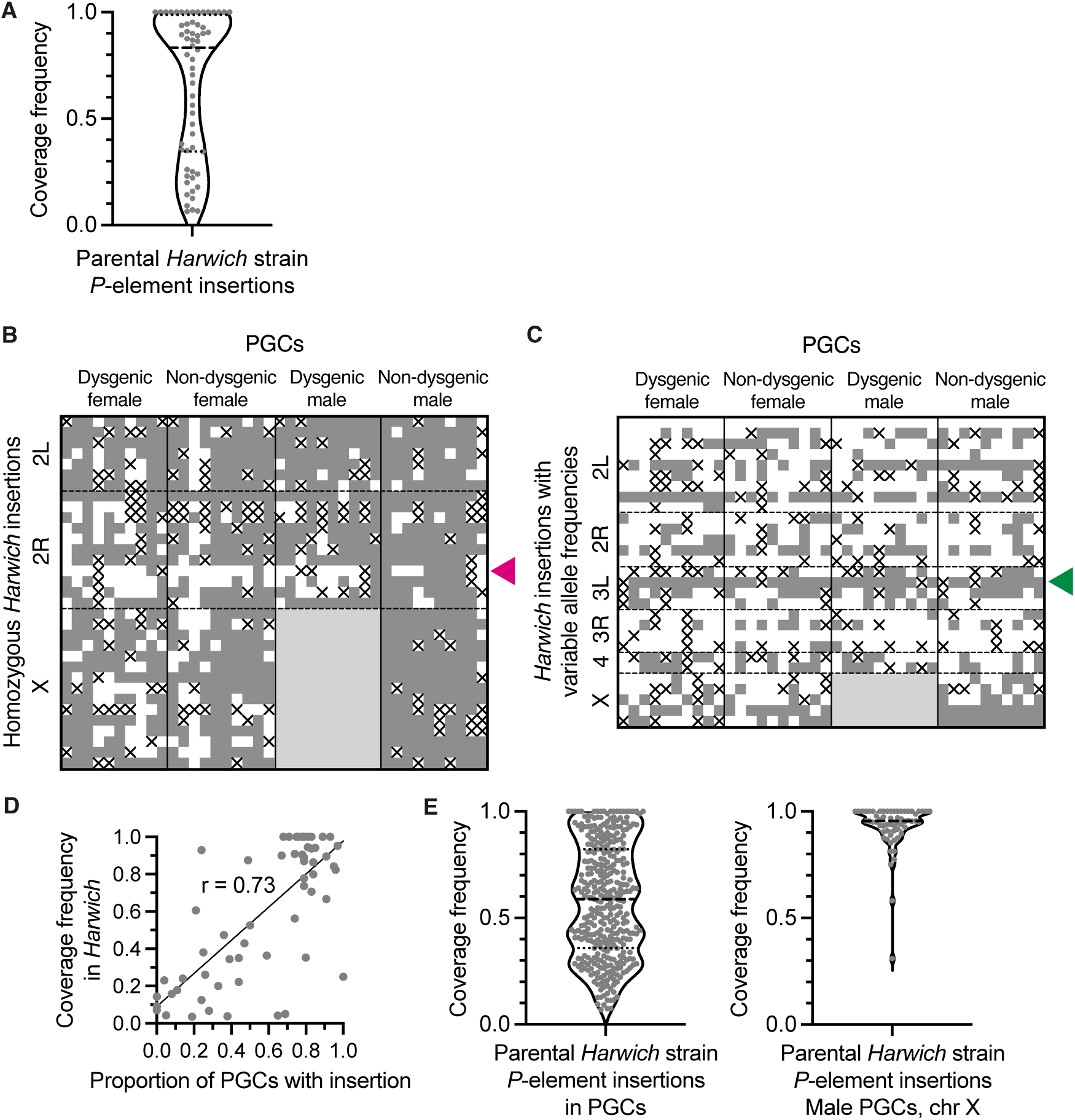
High detection rate of vertically transmitted *P*-elements in PGCs. **A.** Coverage frequencies representing zygosity of *P*-element insertions (grey data points) in the *Harwich* genome. Based on a minimum allele frequency of 0.7 in the population, insertions were defined as homozygous (coverage frequency ≥0.7) or segregating at variable allele frequencies (coverage frequency <0.7). **B-C.** *P*-element insertions (rows) with homozygous (B) or variable allele frequencies (C) in *Harwich* that were detected (grey squares) or not detected (white squares) in the 40 PGC genomes (columns). Black crosses indicate coverage depth at the locus was insufficient to detect insertion (<2 reads). Light grey shaded area represents the *‘white’* X-chromosome of dysgenic males, which lacks *P*-elements. Magenta arrowhead indicates a locus containing two adjacent insertions on chromosome 2R in *Harwich,* which were rarely detected in PGCs. The *nos-moe::EGFP* transgene, which is flanked by short *P*-element sequences, was misidentified by the detection tool as variably segregating in *Harwich* (green arrowhead). **D.** Discovery rate of *Harwich P*-element insertions in PGCs, expressed as the coverage frequency of insertions in *Harwich* against the proportion of PGCs in which the insertion was detected. Black line represents linear regression. r, Pearson correlation coefficient. **E.** Coverage frequencies of *Harwich* insertions detected in PGCs (left panel). Insertions on the X-chromosome detected in (hemizygous) male non-dysgenic PGCs are shown separately (right panel). Dashed line represents median, dotted lines represent first and third quartiles.

**Figure S4.**
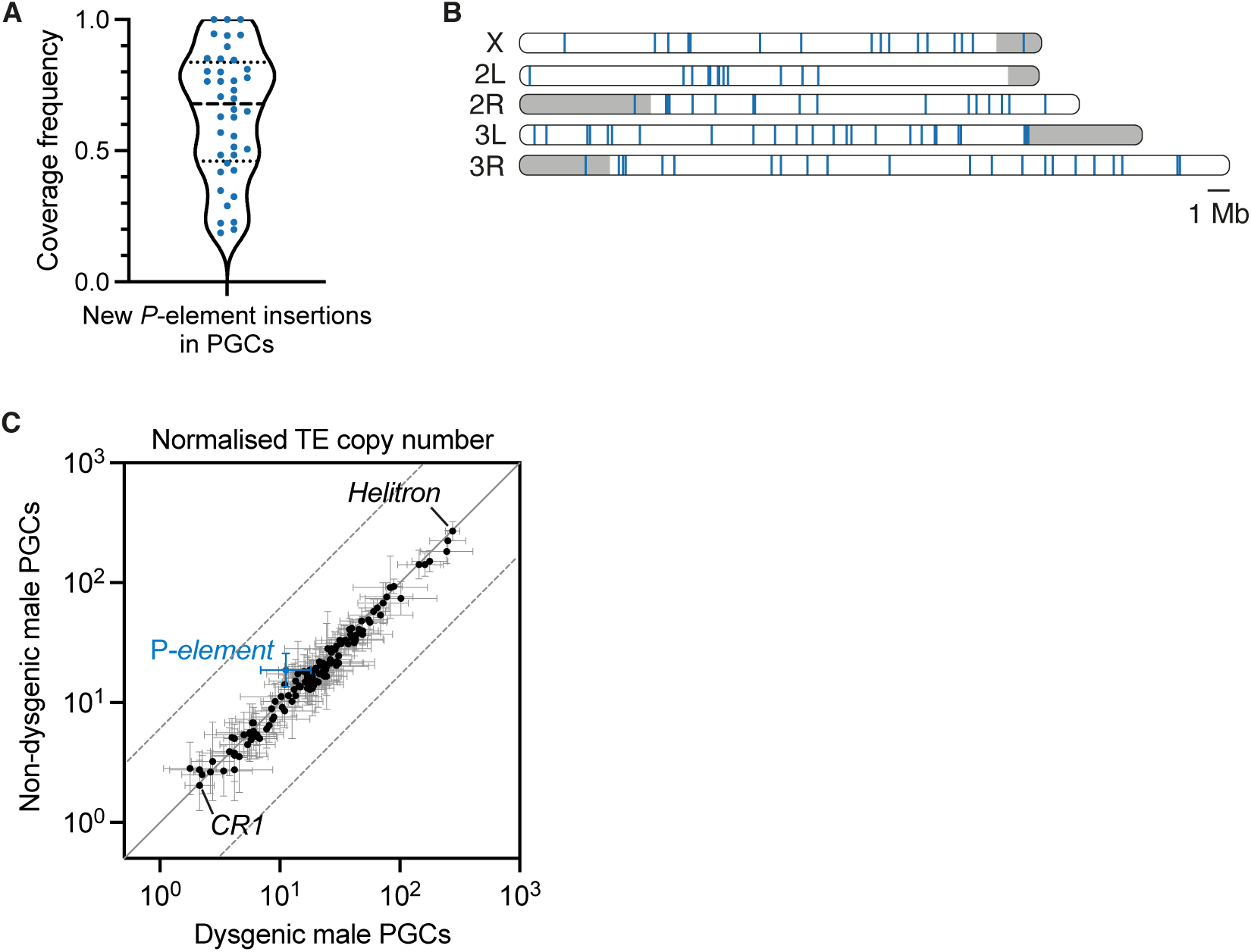
Zygosity and genomic distribution of new *P*-element insertions in PGCs. **A.** Coverage frequencies of new *P*-element insertions (blue data points) detected in PGCs. Dashed line represents median, dotted lines represent first and third quartiles. **B.** Chromosomal distribution of new *P*-element insertions (blue lines) identified in PGCs. Grey shaded regions represent pericentromeric heterochromatic regions. **C.** Scatterplot of normalised genomic copy number for the *P*-element (blue) and 125 other TE families (black) in dysgenic versus non-dysgenic male PGCs. Solid grey line represents perfect correlation. Dashed grey lines indicate 5-fold difference. Error bars are ± one standard deviation.

**Figure S5.**
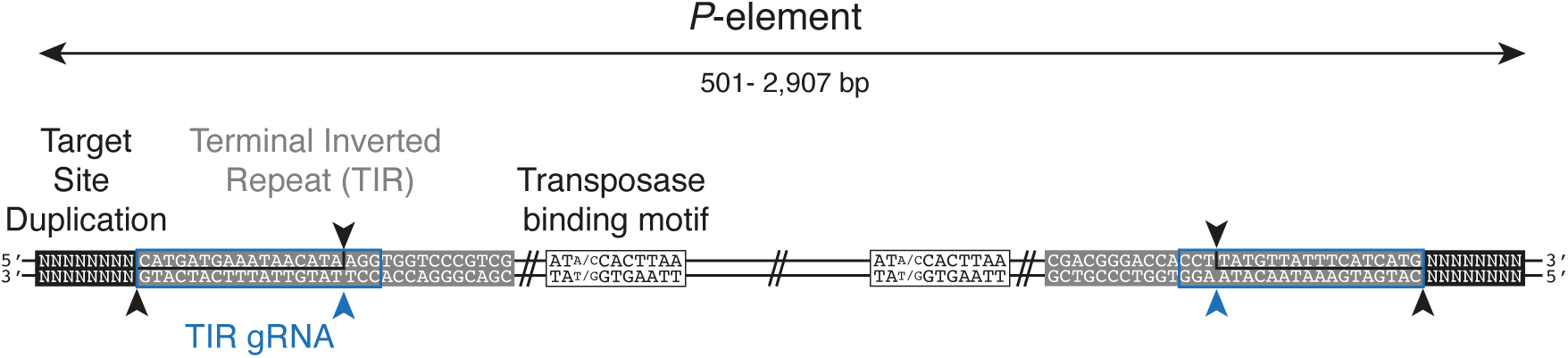
An engineered Cas9 system to induce DSBs at *P*-element TIRs. Schematic showing *P*-transposase cleavage sites (black arrowheads) within 31-bp *P*-element TIRs (grey boxes) at the 5’ and 3’ end of internally deleted and full-length *P*-elements. The TIR-gRNA sequence (blue box) is followed by a PAM (TGG). Cas9 induces DSBs 3 bp upstream of the PAM (blue arrowhead).

**Figure S6.**
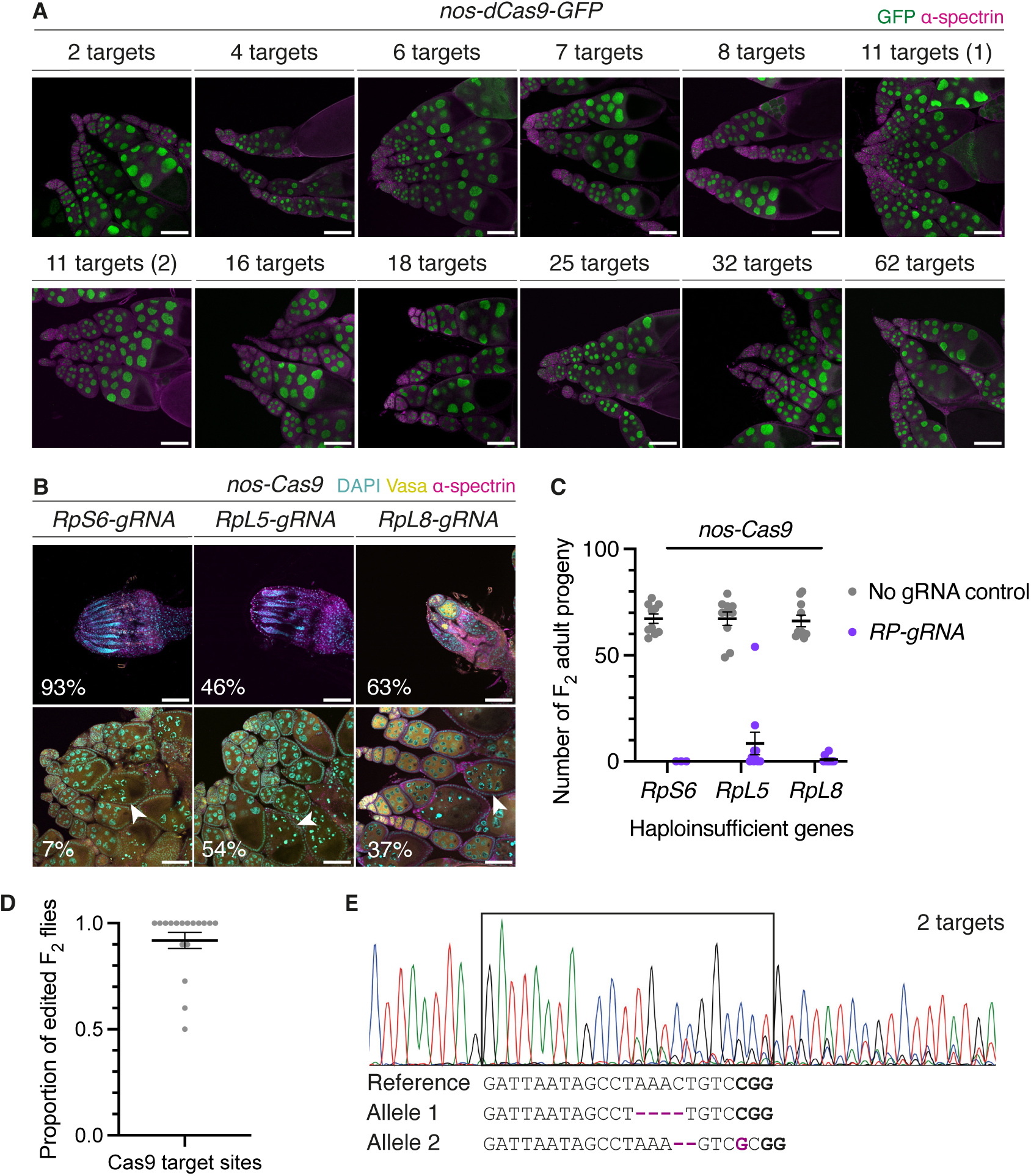
**High efficiency of DSB formation at Cas9 target sites in the germline. A**. Ovaries of F1 progeny from crosses between gRNA-expressing females and males expressing a GFP-tagged Cas9 variant that binds but does not cleave DNA in the germline (*nos-dCas9*), labelled with GFP (green) and α-spectrin (magenta). **B.** Ovaries of F1 progeny from individual crosses between RP-gRNA-expressing females and *nos-Cas9* males, labelled with DAPI (cyan), Vasa (yellow) and α-spectrin (magenta). Percentages represent the relative proportion of progeny with rudimentary ovaries (upper panels) or ovaries with germaria and egg chambers (lower panel). White arrowheads indicate degenerated mid-stage egg chambers with aberrant nuclear morphology. **C.** Fertility tests (n = 10) of adult F1 progeny in (B). Black line is mean, error bars are ± SEM. p < 0.0001 for each pairwise comparison, unpaired *t*-test. **D.** Proportion of F2 individuals (n ≥ 10) carrying 2-, 4-, 6-, 7-, 8- or 11(1)-target gRNA (shown in Figure 4C) harbouring DSB repair products at a given Cas9 target site (grey data points). Black line is mean, error bars are ± SEM. **E.** Sanger sequencing trace showing DSB repair products on the two alleles at the site targeted by 2-target gRNA (relative to the reference sequence). Inserted and deleted bases are shown (magenta). Scale bars, 100 μm.

**Figure S7.**
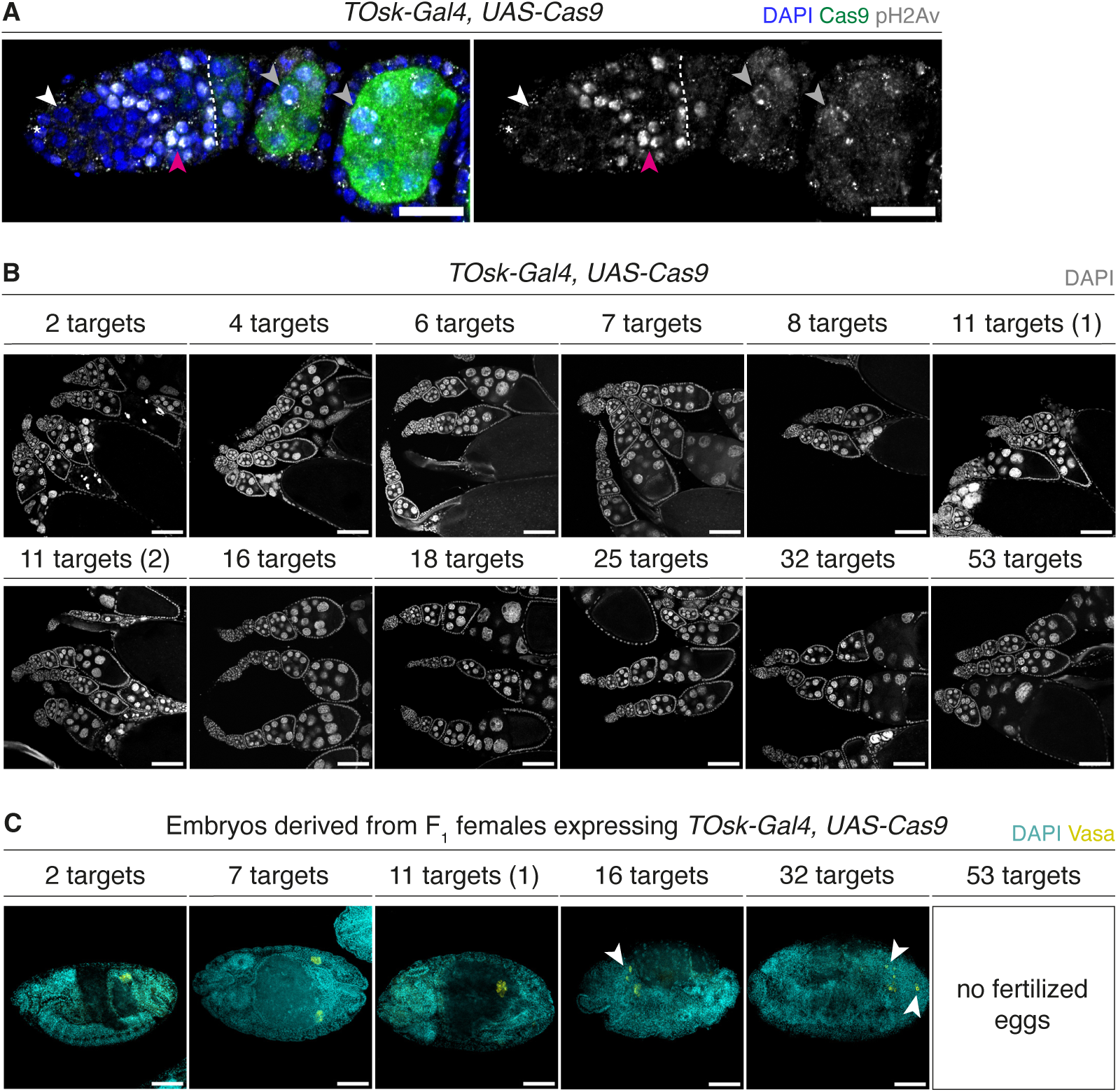
Inducing DSBs in post-mitotic adult germ cells affects embryonic development of the F1. **A.** Germarium from strain expressing Cas9 in the *TOsk* domain (demarcated by dashed line), labelled with DAPI (blue), Cas9 (green) and pH2Av (grayscale). Images are three-channel overlay (left) and single-channel (pH2Av, right). Asterisks indicate GSC niche, white arrowheads indicate GSCs, magenta arrowheads indicate meiotic germ cells and grey arrowheads indicate pH2Av-positive nuclei in early egg chambers. **B.** Adult ovaries of F1 progeny from individual crosses between gRNA-expressing females and *TOsk-Gal4, UAS-Cas9* males, labelled with DAPI (grayscale). **C.** Eggs laid by progeny in (B) aged to ∼16 hours, labelled with DAPI (cyan) and Vasa (yellow). White arrowheads indicate mislocalised PGCs. Scale bars, 20 μm (A) or 100 μm (B and C).

**Table S1.** *P*-element insertions in the *Harwich* strain.

**Table S2.** New *P*-element insertions detected in dysgenic and non-dysgenic hybrid PGCs.

**Table S3.** Copy numbers of *P*-elements in wild-derived and transgenic strains.

**Table S4.** Genomic copy numbers of gRNA sequences within the *nos-Cas9* strain and strains used to generate transgenic gRNA lines (*nos-int;attP2* and *w;TM2/TM6*).

